# Compaction of Active Chromatin Domains and Promoters by H3S10 Phosphorylation during Mitosis Preserves Interphase-Specific Chromatin Structure and Function

**DOI:** 10.1101/2024.05.20.595059

**Authors:** Sweety Meel, Sachin Mishra, Rajat Mann, Arif Hussain Najar, Anurag Kumar Singh, Dimple Notani

## Abstract

Mitotic chromosomes lose interphase-specific genome organization and transcription but gain histone phosphorylation, specifically H3S10p. This phosphorylation event compacts chromosomes in early mitosis by reducing inter-nucleosomal distance before the loading of condensins. However, it is unclear if H3S10p in mitosis preserves the identity of lost chromatin domains and promoters, both physically and functionally. Here, using the pre-mitotic expression of histone H3S10 and its mutants H3S10A and H3S10D, we show that H3S10p hyper-phosphorylates active promoters and spreads into super-domains A in mitosis, causing compaction of these regions. By spreading into active domains in the absence of genome organization, H3S10p retains their identity physically. Functionally, H3S10p ensures optimal closing of promoters by stabilizing the nucleosomes, thereby protecting them from excess loading of transcription machinery post-mitosis. In the H3S10p phospho-mutants, these chromatin regions fail to condense properly during mitosis. As a result, they exhibit enhanced accessibility and transcription of active genes in the next interphase. We propose that the spreading of mitotic H3S10p into active domains preserves their identity during mitosis and, in subsequent interphase, acts as a rheostat to fine-tune transcription and chromatin domain re-formation.

## Introduction

The spatial and temporal regulation of gene expression is crucial in maintaining cell and tissue-type identity. Specific histone modifications, transcription factor binding, and genome organization decorate a given chromatin region for its transcription regulation in interphase. Chromosomes have stable interphase 3D chromatin structures with several degree-folding of the genome, ranging from small-scale fine regulatory loops to megabase-sized TADs (Topologically associating domains) to even larger transcriptionally active (A) and inactive (B) compartments (domains) (Dixon *et al*., 2012; Rao *et al*., 2014; Sexton *et al.,* 2012). During mitosis, these regions undergo hyper-condensation and lose these chromatin features, but the chromatin organisation reappears around late telophase and is revived fully by early-mid G1 (Naumova *et al*., 2013; Nagano *et al*., 2017; Gibcus *et al*., 2018; Abramo *et al*., 2019; Kang *et al*., 2020). How loops, TADs and domains re-emerge post-mitosis is still being determined.

Contrary to the complete loss of chromatin domains, some histone modifications, chromatin remodelers and transcription factors remain bound during mitosis, known as mitotic bookmarking (Martínez-Balbás *et al*., 1995; Valls, Sánchez-Molina and Martínez-Balbás, 2005; Muramoto *et al*., 2010; Zhao *et al*., 2011; Kadauke *et al*., 2012; Caravaca *et al*., 2013; Hsiung *et al*., 2015; Teves *et al*., 2016; Oomen *et al*., 2019; Kang *et al*., 2020; Pelham-Webb *et al*., 2021; Yu *et al*., 2023; Zhu *et al*., 2023). Mitotic persistence of these bookmarkers is critical for enhancer-promoter function and gene transcription in the next interphase.

However, owing to their narrow and local occupancy on chromatin regions, most epigenetic marks and histone modifications are of meagre significance in propagating the memory of large-scale genome folding structures. Nonetheless, the spreading of repressive histone marks, H3K9me2/3 and H3K27me3, in dense inactive B compartments preserves heterochromatin across generations (Nakayama *et al*., 2001; Grewal and Jia, 2007; Owen, Osmanović and Mirny, 2023). A mechanism preserving active euchromatin domains through spreading of an epigenetic mark similar to heterochromatin remains unknown.

Unlike most marks, H3S10p levels in active chromatin increase tremendously from interphase to mitosis (Zhiteneva *et al*., 2017; Javasky *et al*., 2018). H3S10p is known to compact mitotic chromatin by decreasing the inter-nucleosomal distance (Jain, Janning and Neumann, 2021). On the other hand its active suppression is crucial to maintain H3K9me2 heterochromatin regions in interphase (Chen *et al*., 2018; Peng *et al*., 2018). However, whether H3S10p compacts specific genomic regions in mitosis to regulate gene expression in the next interphase remains unexplored.

In this study, we profiled the genomic occupancy of H3S10p in asynchronous and mitotic HeLa cell populations. We observed that H3S10p increases in mitosis and forms islands on mitotic chromatin. Collating our H3S10p ChIP-seq data with HiC, we report that the increase and spread in mitotic H3S10p appears at active interphase domains (A compartment). Moreover, the mitotic H3S10p levels scale with robust transcription in these domains in interphase. Further, active promoters show the highest H3S10p enrichment and are relatively more compacted in mitosis. To test the functional consequence of lack of mitotic H3S10 phosphorylation on these regions and promoters, we used pre-mitotic expression of H3S10 mutants and observed a loss in total H3 incorporation in mitotic chromatin, resulting in lack of chromosome condensation. This led to a widespread opening of active promoters and therefore, abrupt gene transcription in the subsequent interphase as observed by ATAC-seq and RNA-seq. Our data suggest the functional role H3S10p deposition and resultant compaction of active domains and promoters in the regulation of interphase specific transcription.

## Results

### H3S10p marks chromatin as islands in mitosis

H3S10p is present at basal levels in interphase but increases significantly in mitosis, as observed in the snapshots taken from an asynchronous HeLa population (Fig. 1A). H3S10 phosphorylation starts at pericentromeric DNA, as previously reported (Fig. 1A, blue arrows) (Hendzel et al., 1997). It increases in mitosis until late anaphase and is reduced to basal levels by the end of telophase (Fig. 1A-B). A comparison of H3S10p and CENP-B localisation on DNA showed that H3S10p begins around pericentromeric DNA but spreads to different parts of the chromosome as mitosis progresses further (Fig. 1B). We observed that H3S10p levels peaked in prometaphase, where its distribution was largely non-homogenous. It decorated the periphery of condensed chromosomes in a prometaphase HeLa cells, showing highly intense staining of H3S10p at low intensity regions of DNA (Fig. S1A). The peculiar pattern of H3S10p concerning DNA and centromeres was also consistent in prometaphase mouse embryonic stem cells (mESCs) and MCF-7 cells (Fig. 1C, S1A) (Li *et al*., 2005).

**Figure 1.**
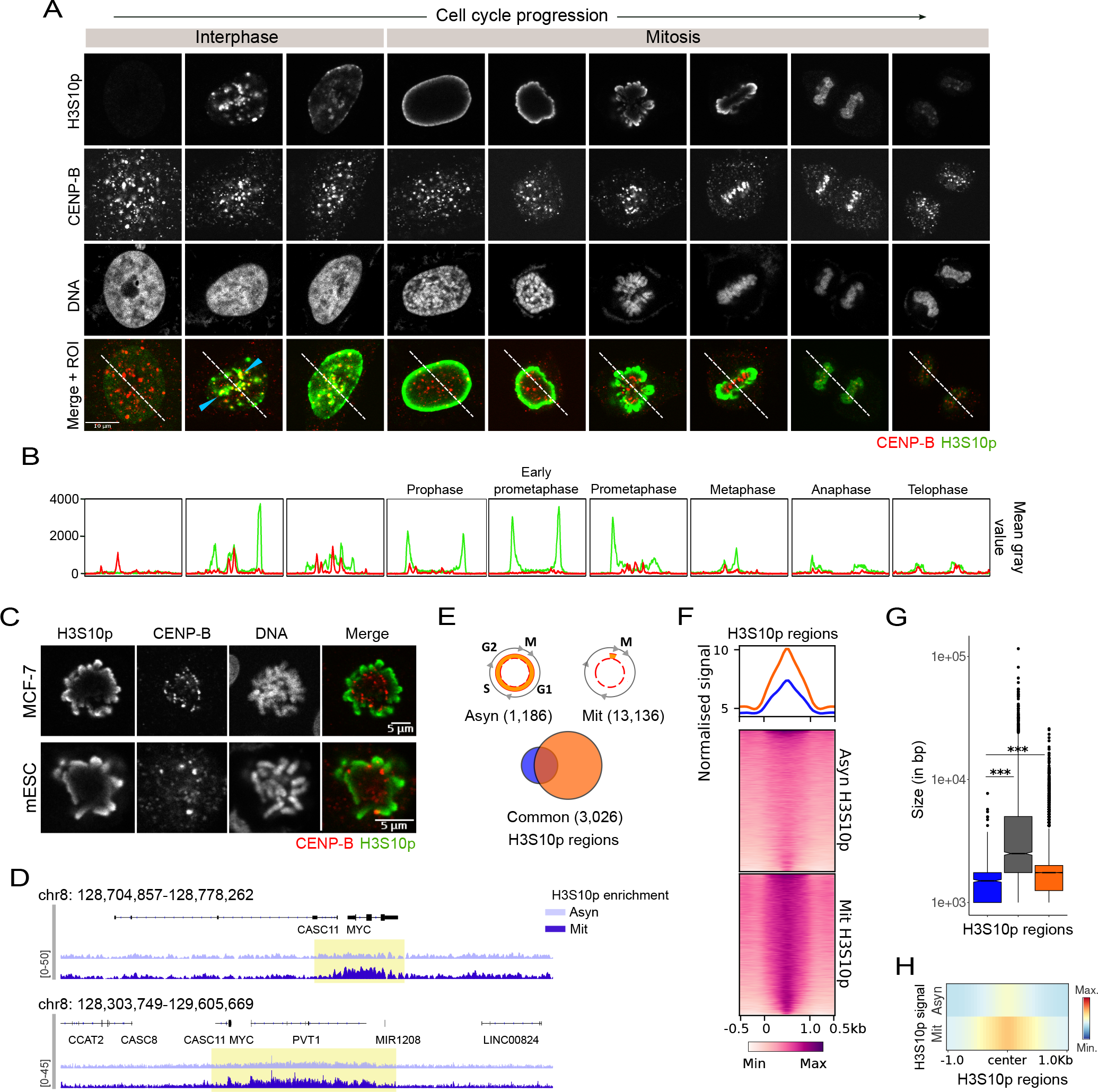
H3S10p marks chromatin as islands in mitosis. A. Snapshots from immunostaining of HeLa asynchronous population showing dynamics of H3S10 phosphorylation during cell cycle. Colours: H3S10p (Green), CENP-B (centromeric protein) in red, and DNA are stained with Hoechst to mark the cell cycle stages. B. Mean grey value plotted for H3S10p and CENP-B along the ROIs from Fig. 1A. C. Snapshot of H3S10p (green) and CENP-B (red) immunostained MCF-7 and mESC mitotic cell to show the prometaphase stage. D. IGV track screenshot of Chr8 showing distinct island-like coverage of H3S10p ChIP-seq in mitotic (Mit, nocodazole treated) but not asynchronous (Asyn, DMSO treated) HeLa population. E. Venn diagram showing the number of H3S10p regions specific and common to the Asyn and Mit population F. Heatmap exhibiting Asyn and Mit H3S10p signal at all H3S10p regions combined (Asyn, Mit, and Common). G. Boxplot showing the size distribution of H3S10p regions from Fig. 1E with an increase in the spreading of H3S10p from asynchronous to mitosis. H. Heatmap profile plot showing the Asyn and Mit H3S10p signal at the H3S10p regions of enrichment. Statistics details: p-value >0.05 (ns), <=0.05 (*), <=0.01 (**), and <0.001 (***). The Mann-Whitney U-test determined statistical significance.

To investigate the specific genomic regions enriched for mitotic H3S10p, we performed ChIP-sequencing of H3S10p in synchronous (prometaphase/mitotic, Mit) and asynchronous (interphase, Asyn) HeLa cells using the well-established nocodazole and DMSO treatment respectively (Fig S1B) (Naumova *et al*., 2013). We observed that H3S10p marks the chromatin in both interphase (Asyn) and mitosis (Mit) (Fig. 1D-F). However, its deposition significantly increased and became largely specific to distinct islands in mitosis (Fig. 1D). Consistent with mitotic H3S10p spreading from foci to larger areas of chromosomes (Fig. 1A), we observed that the size of H3S10p regions was larger in mitosis (Fig. 1G). The highly diffused signal of H3S10p in mitosis indicated that the increase in the size of H3S10p regions was a result of spreading of this modification (Fig. 1F, H). Moreover, the regions common to asynchronous and mitotic populations showed further size expansion in mitosis (Fig. S1E). Our results confirm an increase in H3S10p in mitosis and show that subsequent spreading of H3S10p on DNA gives rise to distinct islands in mitosis. Such spreading of H3S10p as islands might be necessary for mitotic compaction of specific chromatin regions as increased H3S10p levels facilitate chromosome condensation in mitosis (Wei *et al*., 1999; Wilkins *et al*., 2014; Zhiteneva *et al*., 2017).

### Mitotic H3S10p preserves the identity of interphase active domains

We asked if these large mitotic H3S10p marked islands were chromatin domains that decorate interphase chromosomes. Towards this, we compared HiC data from HeLa asynchronous (interphase) population (Rao *et al*., 2014) with mitotic H3S10p ChIP-seq. The analysis revealed many-fold enrichment of H3S10p, specifically in interphase A compartments but not in B compartments (Fig. 2A). Further, the spreading of mitotic H3S10p was restricted to compartment A (Fig. 2B, S2B); therefore, we focused our further analysis on these compartments. We divided the A domains (compartments) into three equal groups (∼500 in each group) based on the mitotic H3S10p levels in them (High, Medium and Low: Mit H3S10p)(Fig. S2C). The compartmentalization score was proportional to the level of mitotic H3S10p in them (Fig. 2C). Since, the deacetylation of histones precedes their phosphorylation, we tested the acetylation status of H3S10p-marked compartments in interphase (Li et al., 2006). Notably, the H3K27ac and H3K9ac deposition was high in these compartments (Fig. 2D, S2A) (Calo and Wysocka, 2013; Beacon *et al*., 2021; Zheng *et al*., 2022). This correlation was also valid for their interphase transcription (Fig. 2E). A domains with higher H3S10p deposition also showed higher gene density and a more significant proportion of these genes expressed in interphase (Fig. S2D-E). A domains are an aggregate of active TADs and therefore, indeed, TADs recapitulated the trend observed for compartments/domains. Furthermore, we found that the TADs in high H3S10p-marked A domains had robust boundary insulation and higher CTCF, Rad21, and SMC3 occupancy at their boundaries (Fig. 2F-G, S2F).

**Figure 2.**
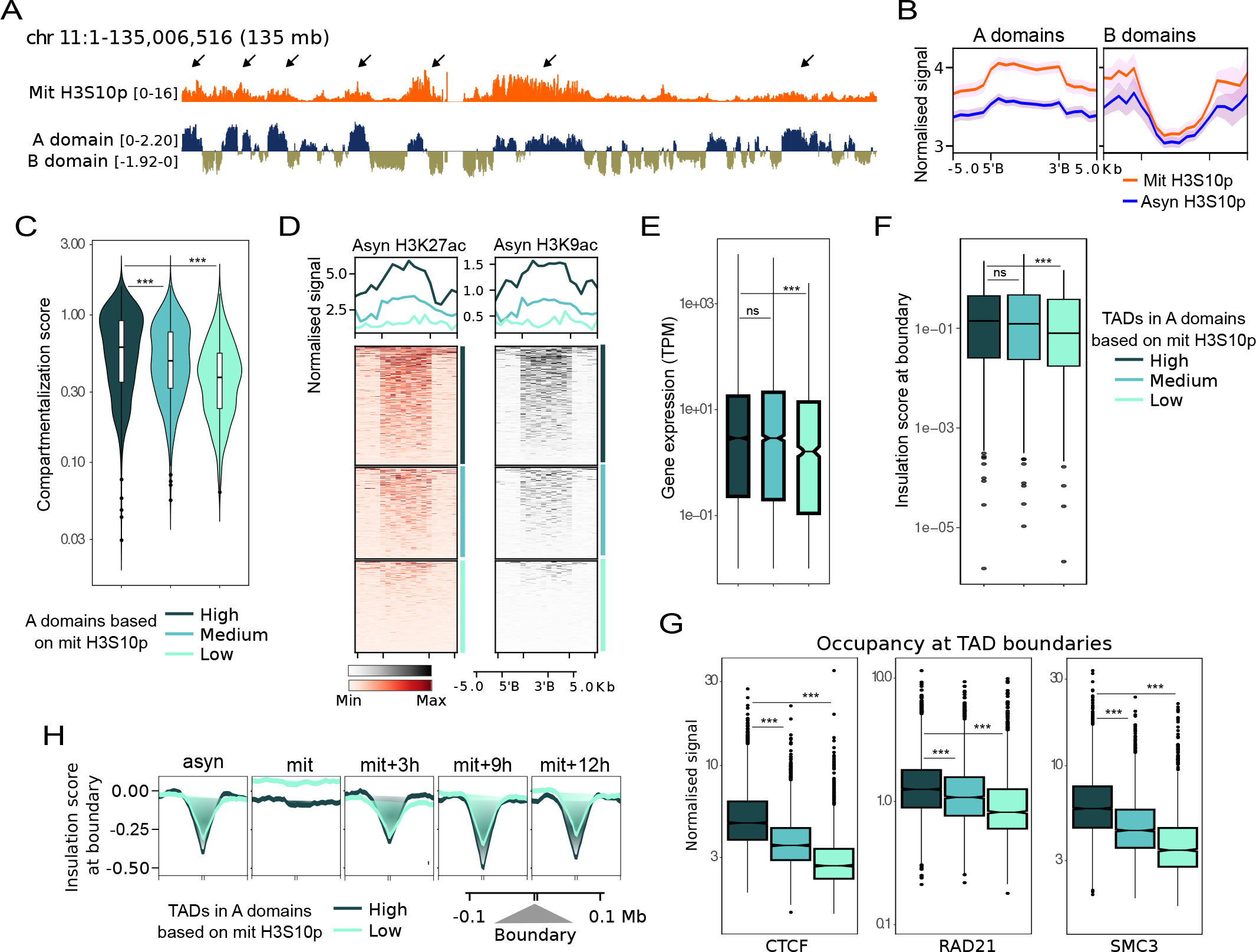
Mitotic H3S10p preserves the identity of interphase active domains. A. IGV track screenshot of Chr 11 showing A (active) and B (inactive) compartments/domains (HiC) with mitotic H3S10p ChIP-seq track. B. Profile plot showing Asyn and Mit H3S10p signal at interphase A and B domains. C. Violin plot showing compartmentalization strength of A domains categorized into three groups based on mitotic H3S10p levels. D. Heatmap with interphase H3K27ac signal at the three categories of A domains. E. Boxplot with gene expression (interphase RNA-seq, TPM) of genes present in the three A domain categories. F. Boxplot with the distribution of insulation score at the TAD boundaries of the TADs found in the three A domain categories. G. Boxplots showing interphase CTCF, RAD21, and SMC3 occupancy at the TAD boundaries of the three A domain categories. H. Profile plot showing insulation score at TAD boundaries of TADs with low and high H3S10p enrichment indicating boundary strength at different time points post-mitosis. Statistics details: p-value >0.05 (ns), <=0.05 (*), <=0.01 (**), and <0.001 (***). The Mann-Whitney U-test determined statistical significance.

TADs are known to re-emerge around the end of telophase during mitosis and acquire stable structure by mid-G1 (Dileep *et al*., 2015; Abramo *et al*., 2019; Kang *et al*., 2020; Pelham-Webb *et al*., 2021). We asked if the levels of mitotic H3S10p in TADs were correlated with their re-appearance post mitosis. Analyzing publicly available HeLa HiC data at various time points from mitotic release (Abramo *et al*., 2019) revealed the early post-mitotic onset of TADs with high mitotic H3S10p enrichment, with better insulation throughout the cell cycle (Fig. S2G, 2H). These TADs exhibited high intra-TAD interactions throughout the cell cycle compared to high H3K27ac TADs (Fig. S2H). Together, we conclude that in the absence of interphase-specific chromatin organization in mitosis, mitotic H3S10p spreading physically marks the interphase A compartments/domains as per their transcriptional activity in interphase.

### H3S10p prefers promoters over enhancers in mitosis

We observed that within the H3S10p-marked islands, specific genomic regions showed more localised H3S10p enrichment (Fig. 3A). Genome-wide annotation of these regions identified them primarily as promoters (Fig. 3B). The onset of mitosis is accompanied by global deacetylation including the loss of H3K27ac, an active chromatin mark (Valls, Sánchez-Molina and Martínez-Balbás, 2005; Javasky *et al*., 2018; Kang *et al*., 2020). Comparison of H3S10p and H3K27ac from asynchronous and mitotic conditions exhibited a higher loss of H3K27ac at promoters and a corresponding higher gain of H3S10p in mitosis, with relatively minor differences at enhancers (Fig. 3C, S3A). Together, this data suggests promoter bias in mitotic H3S10p enrichment.

**Figure 3.**
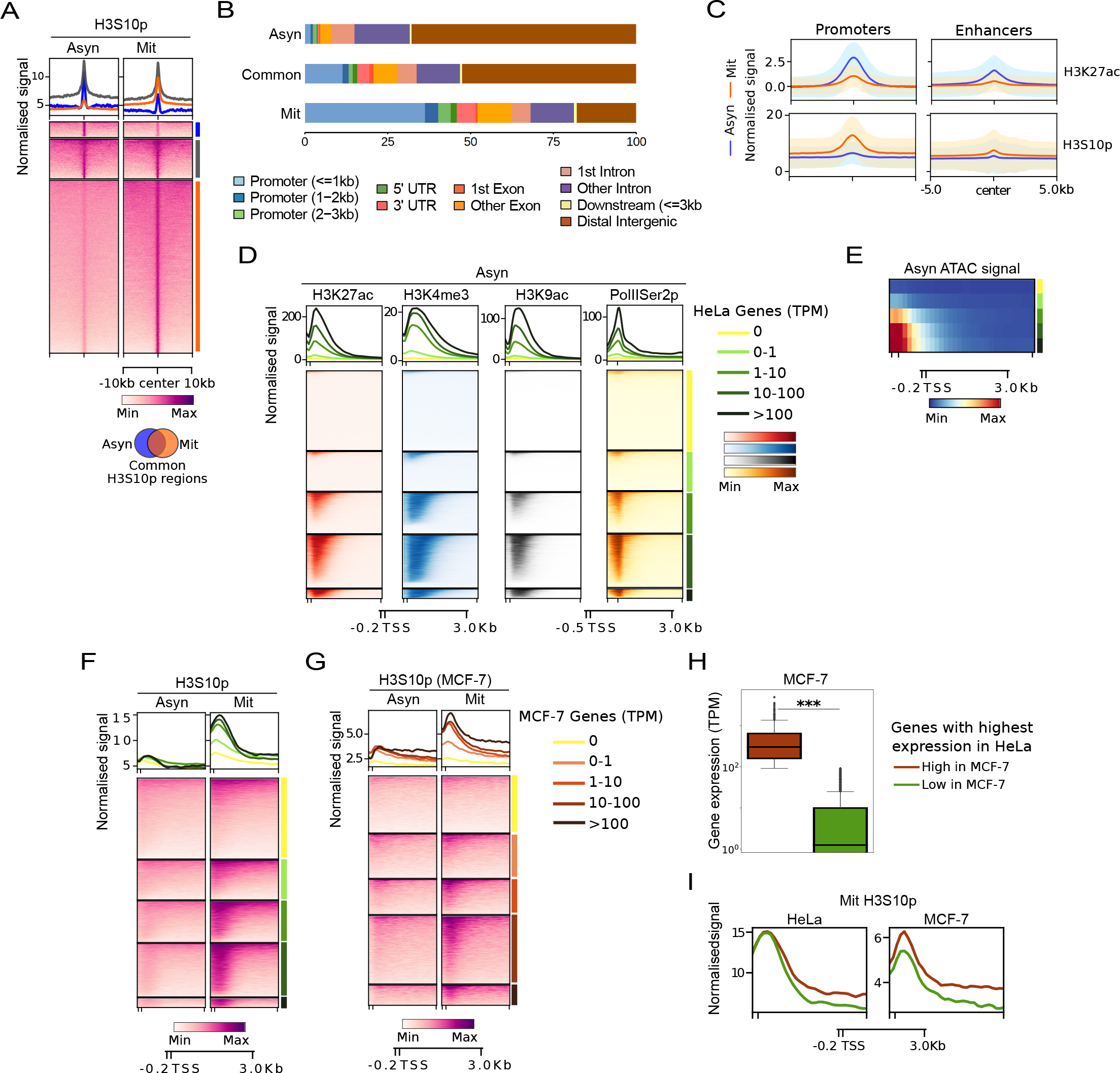
H3S10p prefers promoters over enhancers in mitosis. A. Heatmap showing Asyn and Mit H3S10p signal at asynchronous and mitosis specific (Asyn and Mit), and regions common to both (Common). B. ChIP seeker analysis of Asyn, Mit, and Common H3S10p regions reveals greater promoter enrichment of H3S10p in mitosis. C. Profile plot with H3K27ac and H3S10p (mitotic and asynchronous) signal at HeLa enhancers (H3K27ac peaks without TSS) and promoters (H3K27ac peaks with TSS). D. Heatmaps showing levels of active histone marks (H3K27ac and H3K4me3) and elongating Pol-II (Pol-IIS2p) from asynchronous populations at the gene categories binned based on HeLa gene expression (TPM). E. Profile heatmap showing ATAC signal from asynchronous and mitotic populations at the gene categories described in Fig. 3D. F. Heatmap showing Asyn and Mit H3S10p signal at promoters of genes categorized based on HeLa gene expression (TPM). G. Heatmap showing Asyn and Mit H3S10p signal from MCF-7 cells at promoters based on MCF-7 gene expression (TPM). H. Gene expression in MCF-7 cells from the two subgroups of genes with highest expression in HeLa having high or low expression in MCF-7 I. Mitotic H3S10p signal from HeLa and MCF-7 cells at the two subgroups of genes with highest expression in HeLa having high or low expression in MCF-7. Statistics details: p-value >0.05 (ns), <=0.05 (*), <=0.01 (**), and <0.001 (***). The Mann-Whitney U-test determined statistical significance.

To test if high levels of H3S10p on promoters in mitosis have any association with the gene transcription in interphase, we binned the promoters into four transcribing and one non-transcribing category in HeLa. Respective enrichment of active histone marks (H3K4me3, H3K9ac, H3K27ac), elongating RNA polymerase (PolIISer2p), and chromatin accessibility in interphase validated the transcriptional binning of promoters (Fig. 3D-E). Interestingly, mitotic H3S10p levels at promoters scaled with the transcriptional activity of genes in interphase (Fig. 3F, Right panel). On the other hand, the asynchronous H3S10p signal on these promoters did not show significant differences (Fig. 3F, left panel). These results suggest that within the islands of mitotic H3S10p, active promoters are hyper-marked by H3S10p in mitosis. Further, this marking is proportional to their transcription in interphase. This is consistent with the recent studies describing prioritized mitotic bookmarking of high-expressing gene promoters over regulatory enhancers (Yu *et al*., 2023; Zhu *et al*., 2023).

To test if promoter marking by H3S10p is cell-type specific, we performed H3S10p ChIP-seq in mitotic and asynchronous MCF-7 cells. Similar to the trend observed at promoters binned based on transcription in HeLa cells, we observed H3S10p enrichment scaled to gene expression upon promoter binning in MCF-7 (Fig. 3G). For assessing cell type specific promoter enrichment of mitotic H3S10p, we took genes from the highest expression quantile in HeLa cells (∼2000 genes). The expression of these genes varied over a wide range in MCF-7, so we divided them into high (∼1100) and low (∼900) expressing subgroups in MCF- 7 (Fig. 3H). The H3K27ac levels at the promoters corroborated with their expression in a cell-type specific manner (Fig. S3B). Supporting our hypothesis, the mitotic H3S10p levels also differed concordantly at the two subgroups in MCF-7 while HeLa showed no difference (Fig 3I). Our data suggest that mitotic H3S10p marks the promoters in a transcription dependent manner. Additionally the marking of active promoters by H3S10p is cell-type specific, scaling to their transcriptional potential.

### Lack of H3S10 phosphorylation in mitosis results in reduced compaction of active domains

Our observations so far have established a positive correlation between mitotic H3S10p deposition and interphase chromatin structure and function. Next we asked if H3S10p has any causative functional role in chromosome compaction and if the defects in H3S10p mediated compaction is important in ‘maintaining’ the state of interphase chromatin. Towards this, we decided to create phospho-mutant lines. Creating in-vivo mutations in histone genes is technically challenging as they exist in gene clusters. Notably, the mutation at H3K27 (Lysine) to M (Methionine), only in a few gene copies causes paediatric Diffuse Midline Glioma (DMG) (Chan *et al*., 2013; Sahu and Lu, 2022). Hence, we made three HeLa stable cell lines expressing either of the three inducibly; a) wild type (H3S10, Serine), b) phospho-dead mutant (H3S10A, Alanine), c) phospho-mimetic mutant (H3S10D, Aspartic acid) (Fig. S4B-C). These exogenous histones with HA-peptide fusion were doxycycline inducible (Fig. 4A). We confirmed the incorporation of HA-tagged histones into mitotic chromatin by HA-Immunoprecipitation (Fig. S4D). in order to understand the function of mitotic H3S10 phosphorylation *per se* and to avoid interphase-borne effects, we optimised the doxycycline treatment time window adequate enough for pre-mitotic expression in S-G2 stage, followed by mitotic arrest (Fig. 4B). Upon induction, the gradual increase in HA-tagged histone incorporation in chromosomes was confirmed (S5A-C). We only allowed low level of induction to abstain from alterations in the stoichiometry of total histone H3 (Fig. S5D).

**Figure 4.**
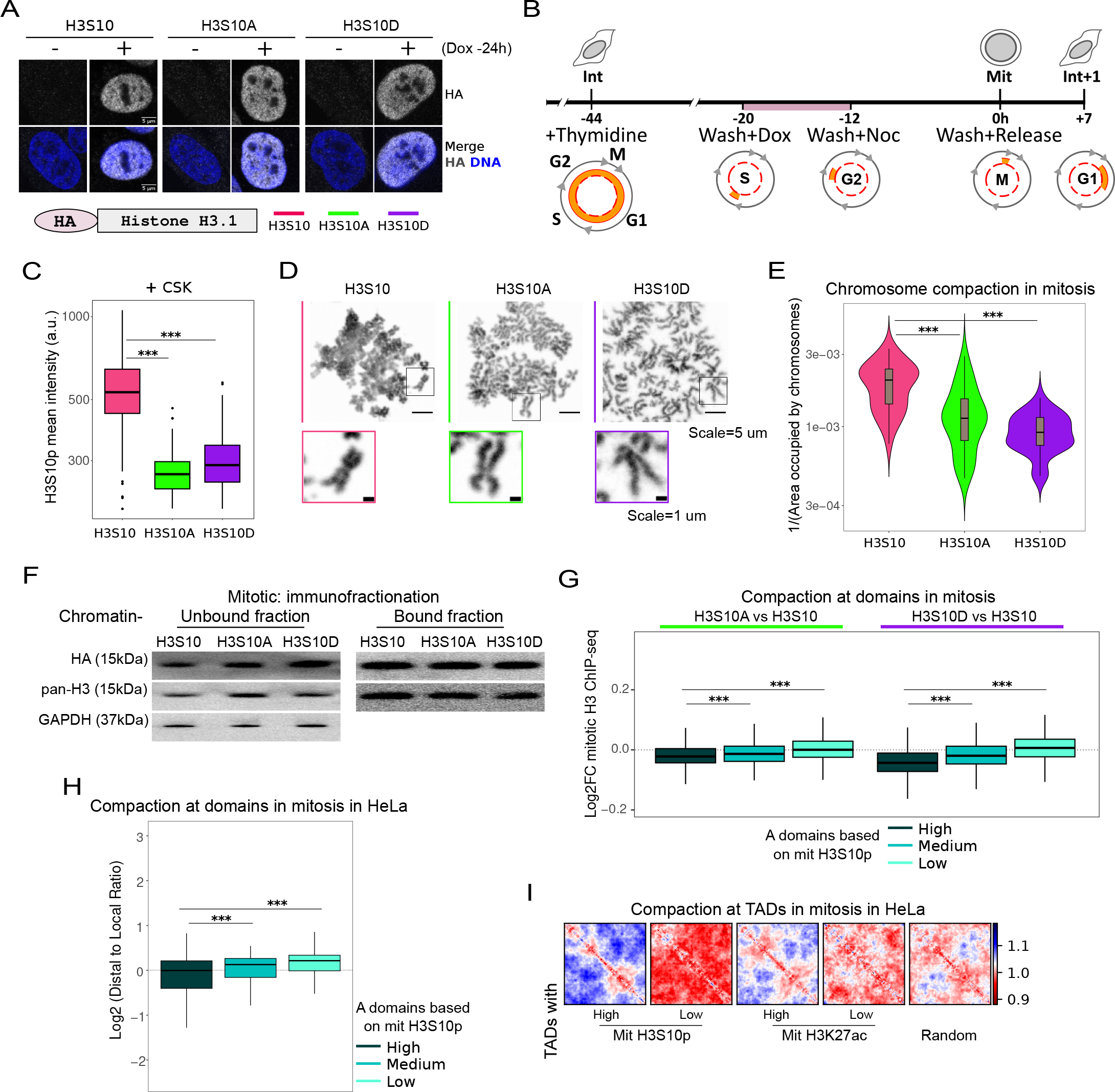
Lack of H3S10 phosphorylation in mitosis results in reduced compaction of active domains. A. HA immunostaining upon doxycycline induction in stable lines expressing: HA-H3S10 (wild-type), HA-H3S10A (phospho-dead mutant), and HA-H3S10D (phospho-mimetic mutant). The panel below shows the schematic of HA-H3S10 and mutants. B. Schematic of the treatment regime followed for mitosis-specific induction of HA-tagged histone H3 variants and release into next interphase. C. Boxplot showing chromatin-bound H3S10p immunostaining of mitotic cells treated with CSK buffer. D. Metaphase spread of chromosomes from the three HeLa lines. Insets highlighting inconsistent chromosome morphology in H3S10A and H3S10D lines. E. Violin plot showing chromosome compaction from the three HeLa lines; measured as an inverse of the area occupied by a single nucleus metaphase spread. F. Immunoblot of chromatin - unbound and bound H3S10p, HA, and total H3 levels in the three HeLa lines. G. Boxplots showing the change in total H3 levels at active domain categories (from Fig. 2C) in mutant vs wild type line, as a proxy of mitotic chromatin compaction. H. Boxplots showing actual mitotic chromatin compaction measured as Log2DLR from mitotic HiC, at active domain categories (from Fig. 2C) in HeLa. I. APA plot depicting mitotic compaction at various TAD categories in mitosis. Statistics details: p-value >0.05 (ns), <=0.05 (*), <=0.01 (**), and <0.001 (***). The Mann-Whitney U-test determined statistical significance.

The lines showed no difference in HA levels in mitosis upon immunostaining suggesting equal incorporation of HA-tagged histones (Fig. S5E-F). To examine if mutants showed less H3S10p levels in mitotic chromatin, we performed immunostaining of mitotic cells with CSK (Cytoskeletal) buffer treatment to remove any soluble pool of histones. Indeed, we observed loss of H3S10p in H3S10A and H3S10D mutant lines (Fig. 4C). Since H3S10p facilitates condensation of mitotic chromosomes, the mutant lines showed elongated and thinner chromosomes as opposed to those from wild type line (Fig. 4D). To quantify these features further, we plotted the inverse of the area occupied by the mitotic chromosomes as a measure of their compaction and found that the mitotic chromosomes in mutants had significantly reduced compaction (Fig. 4E). The lack of condensation can be a result of increased nucleosomal eviction or stabilization, so we tested the total H3 levels as proxy for nucleosome numbers in mitotic chromatin fraction between the lines. (Fig. S5G). Indeed, both the mutants exhibited reduced presence of chromatin-bound total H3 (Fig. 4F). This observation was further confirmed by the ChIP-seq of H3 that also showed loss of signal in mutants, specifically in A domains that otherwise show high H3S10p enrichment (Fig. 4G). The data suggested that loss of H3 in mutants was proportional to gain of mitotic H3S10p in WT. We hypothesised that H3S10p’s presence potentially stabilises the nucleosomes for mitotic chromatin compaction. Indeed, mitotic HiC data revealed that the active A domains and TADs were compacted in proportion to their mitotic H3S10p levels (Fig. H-I, S5H). We did not observe similar trend for the control TADs with varying mitotic H3K27ac levels indicating that higher mitotic compaction in wild-type and greater loss of H3 in mutants, was specific to H3S10p enriched TADs (Fig. I, S5H). Our results show that stabilisation of nucleosomes by H3S10p is required for compaction of active domains in mitosis that are relatively more open and transcribe in interphase.

### Active genes are further up-regulated in the next interphase upon loss of mitotic compaction at promoters

H3S10p enrichment in mitosis was the highest at active promoters scaling to their interphase transcriptional activity (Fig. 3F). Given high H3S10p levels in active domains/TADs conferred more compaction in mitosis (Fig. 4H-I), we asked if high H3S10p levels at promoters were related to their mitotic compaction. Intriguingly, most active promoters showed maximum loss of ATAC-signal in mitosis, potentially to safe guard them from spurious transcription activity in mitosis (Fig. 5A, upper panel). Aurora Kinase B (AURKB) catalyses mitotic H3S10 phosphorylation preferably at hypoacetylated H3K9ac histones (Li et al., 2006). As a result, we observed a combined loss and gain of H3K9ac and H3S10p respectively at the promoters in mitosis; in a transcription scaled manner (Fig. 5A, lower panel). We speculate that the yin-yang between these two histone modifications might be crucial for the loss of ATAC-signal seen at those respective promoters in mitosis for their optimal compaction.

**Figure 5.**
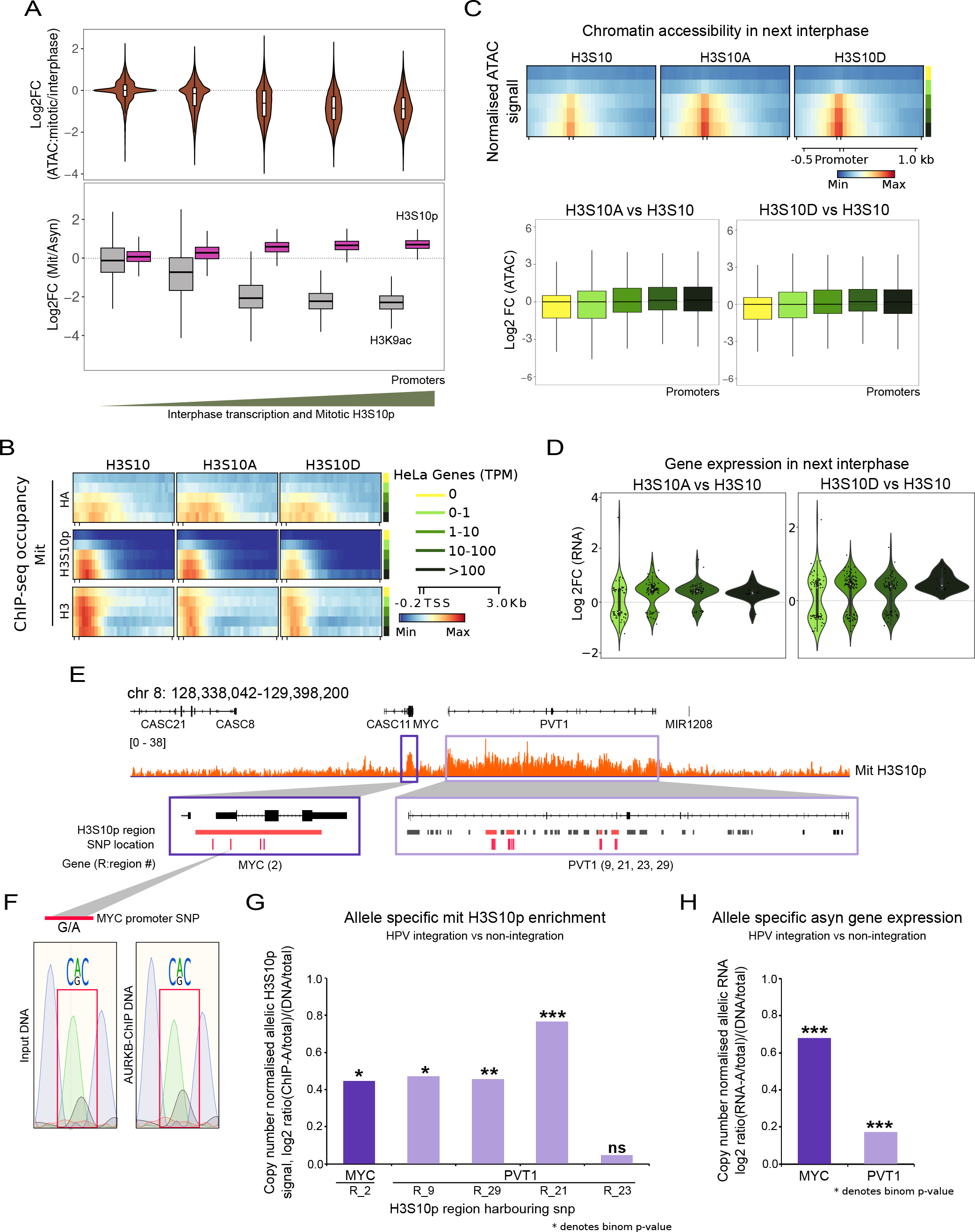
Active genes are further up-regulated in the next interphase upon loss of mitotic compaction at promoters. A. Violin plot showing loss of ATAC from interphase to mitosis (upper), and box plot showing loss of H3K9ac and gain of H3S10p from interphase to mitosis (lower) at promoter categories from Fig. 3D. B. ChIP-seq profile of mitotic HA, H3S10p, and total H3 from the three stable lines at the promoter quantiles in HeLa from Fig. 3D. C. Profile plot (upper) and box plot (lower) showing the increased opening in the next interphase at promoter categories from Fig. 3D in H3S10, H3S10A and H3S10D lines. D. Violin plots showing RNA-seq fold changes in H3S10A (left panel) and H3S10D (right panel) lines as a ratio of gene expression compared to H3S10. The changes were plotted at the gene categories in Fig. 3D. E. IGV track screenshot of chromosome 8q24 locus showing mitotic H3S10p enrichment at MYC and PVT1 genes with multiple allele-specific SNPs. F. Sanger sequencing of MYC promoter amplified from the mitotic-HeLa input and AURKB-ChIP DNA showing allelic difference at the respective SNP. G. Bar plot showing the H3S10p levels at various regions with SNPs in MYC and VT1 gene from the HeLa Haplotype A with HPV integration. H. Bar plot showing the MYC and PVT1 gene expression levels from the HeLa Haplotype A with HPV integration. (G-H) The y-axis represents the log2 ratio of the {proportion of (RNA/ChIP) reads from Haplotype A out of the total (RNA/ChIP) reads} by the {proportion of DNA reads from Haplotype A out of the total DNA reads}, at heterozygous SNPs in each of the features. A binomial test was done for the RNA/ChIP reads from Haplotype A against the proportion of DNA reads from Haplotype A, at heterozygous SNPs in each of the features. Statistics details: p-value >0.05 (ns), <=0.05 (*), <=0.01 (**), and <0.001 (***). The Mann-Whitney U-test determined statistical significance.

In order to understand if H3S10p and H3 loss in mutant mitotic chromosome was co-related to their transcriptional potential we performed ChIP-seq for H3S10p and HA in WT and mutant lines in mitosis. Notably, the HA levels were similar between the wild type and mutant lines suggesting equal incorporation of exogenous histones into the chromatin (Fig. 5B).

However, the most active promoters showed a greater loss of H3S10p as well as total H3, same as domains (Fig. 4G), in both the mutant lines (Fig. 5B). The data suggest reduction in local mitotic compaction at the active promoters; which in turn might hamper the mitotic transcriptional shut down of genes.

Next, we asked how the loss of chromatin compaction localised at promoters was functionally relevant to their transcriptional regulation in the next interphase. For this, we harvested cells at different time points post-mitosis (Fig. S7A). We chose the time point of 7 hours for further experiments as the stable chromatin interactions and transcription resume by early-mid G1, between 6-8 hours in mammalian cells (Abramo *et al*., 2019; Kang *et al*., 2020). We first confirmed negligible levels of exogenous histones in the next interphase by HA immunoblotting (Fig. S7B), indicating that any effects observed in this stage would be a consequence of histone expression in mitosis. Furthermore, wild-type or mutant histones in chromosomes would be diluted away as canonical H3 is largely replaced by H3.3 in interphase during transcription activation.

We then performed ATAC-seq and RNA-seq in the next interphase (7 hours post-mitosis) following 8 hours of pre-mitotic induction of histone variants as described earlier (Fig. 4B). We observed increased chromatin-accessibility at promoters in the next interphase in the mutants, while enhancers remained minimally affected (Fig. S7C). The most active promoters showed highest levels of H3S10p in mitosis (Fig. 3F) and exhibited respective greater loss of H3 among all the promoters (Fig. 5B), suggesting their maximum dependence on H3S10p for compaction in mitosis and probably also their transcription in the next interphase. Indeed, when binned based on transcription, genes with highest expression showed hyper-promoter accessibility in the next interphase in the mutants (Fig. 5C). When compared with RNA-seq, alterations in promoter opening led to further up- regulation of respective genes in the next interphase (Fig. 5D). Therefore, the data suggest that high levels of mitotic H3S10p at active promoters ensures their adequate closing in mitosis. The lack of promoter closing in the mutants results in carrying over of an additional quotient of chromatin openness and transcriptional potential in the next interphase.

H3S10 phosphorylation in mitosis is majorly catalysed by AURKB (Komar and Juszczynski, 2020; Prigent and Dimitrov, 2003). Despite forming islands on active domains, H3S10p enrichment is highest at promoters in mitosis. Hence, we hypothesized that AURKB is loaded on promoters; in a transcription dependent manner. To test this hypothesis, we chose 8q24 locus in HeLa which is heterozygous for HPV integration (Fig. 5E). The *Myc* and *PVT1* genes from Haplotype A (integrated allele) and Haplotype B (non-integrated allele) show differences in interphase transcription as measured from multiple phased SNPs in these genes (Singh *et al*., 2022). *Myc* has “G to A” SNP in its promoter in Haplotype A and we found that the ratio of A:G dominance was maintained in AURKB ChIP-IP DNA similar to that in input DNA (Fig. 5F). This shows that AURKB preferably binds to the *Myc* allele with HPV integration. SNP-based read segregation of H3S10p regions at *Myc* promoter also revealed a significantly enhanced enrichment of mitotic H3S10p at Haplotype A (Fig. 5G). Further, the Haplotype A contributed to majority of *Myc* transcription in interphase (Fig. 5H). A similar trend was observed for *PVT1* gene on Haplotype A (Fig. SG-H). Taken together, these observations confirm specific loading of AURKB on transcriptionally active promoters.

Our study suggests a specific and targeted enrichment of mitotic H3S10p at active chromatin regions based on their interphase transcriptional status. High H3S10p levels allow adequate compaction of these regions in mitosis (relatively the most open regions in interphase), which is crucial for their fine-tuned transcriptional reactivation post-mitosis.

## Discussion

In this study, a genome-wide investigation of H3S10p enrichment reveals that this histone modification spreads into active domains (A compartments) for their compaction in mitosis (Fig. 6A). Within these transcriptionally active domains, promoters exhibit the highest enrichment of mitotic H3S10p scaling to their transcription in interphase. Further, mitotic enrichment of H3S10p at promoters is allele and cell-type specific. Pre-mitotic expression of exogenous H3S10A or H3S10D mutant histones results in enhanced chromatin accessibility and hyper-activation of high transcribing genes in the next interphase (Fig. 6B, upper panel). These interphase defects arise due to the reduced H3S10p levels in mitosis, leading to loss of total H3/nucleosome incorporation into active domains and promoters and hence defective chromosome compaction (Fig. 6B, lower panel). In summary, enhanced H3S10p enrichment at active chromatin domains and promoters ensures their adequate compaction in mitosis which in turn is crucial for the resetting of accurate transcriptional levels in the next interphase. These observations are in line with a recent study showing the importance of mitotic chromosome condensation in the transcription of next interphase (Ramos-Alonso *et al*., 2023).

**Figure 6.**
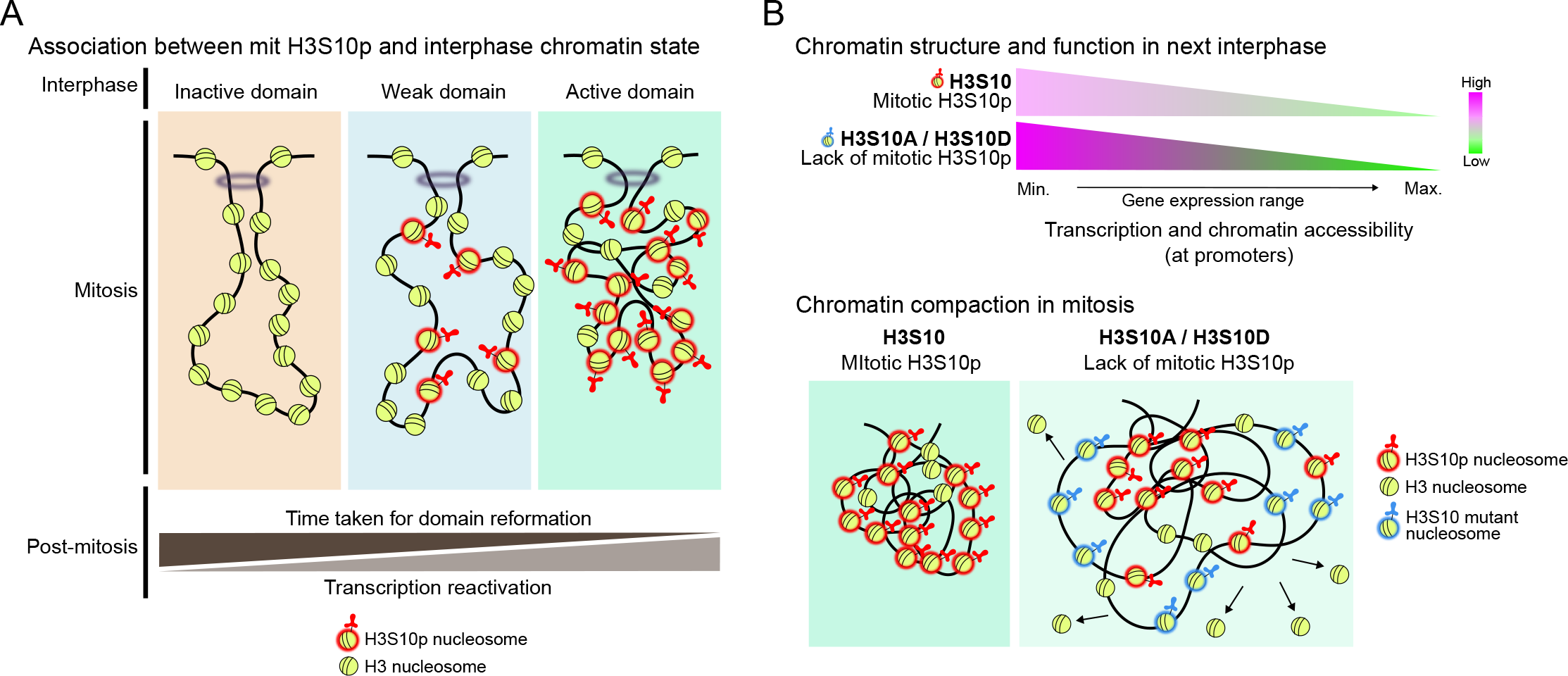
Model. Mitotic chromatin compaction by spreading of H3S10p in active domains stably preserves their interphase-specific 3D structure and transcription. Active domains from interphase have the highest H3S10p levels in mitosis and hence are most compacted relatively. Mitotic proximity of regulatory regions in these domains facilitates their early reformation and robust transcription reactivation post-mitosis (A). Pre-mitotic expression mutant (H3S10A and H3S10D) histones result in hyperactivation of high transcribing genes in the next interphase (B, upper). This is a consequence of a surge in promoter potential in this stage which in turn happens due to the loss of H3S10p and related decrease in chromatin compaction during mitosis (B, lower).

### Active chromatin and promoter condensation by H3S10p

AURKB is recruited on chromatin by HDAC3 (Li et al., 2006). HDAC3 deacetylates H3K9ac which is primarily present on promoters and its levels on promoters are proportional to their transcriptional activity. In this regard, we observe that promoters with high H3K9ac in interphase also exhibit proportional high levels of H3S10p, suggesting, deacetylation of H3K9ac by HDAC3 recruits AURKB for H3S10p deposition at promoters before the onset of mitosis. Further, the transcriptional activity is determined by promoter openness in interphase, but in mitosis a proportional H3S10p enrichment is required to fill the nucleosomal free regions at the active promoters. Indeed, we do observe the positive correlation between promoter opening, transcription and H3S10p deposition. The lack of which, causes loss of nucleosomes at the most active promoters indicating, to condense the promoters; H3S10p stabilises nucleosomes before decreasing the inter-nucleosomal distance. Loss of total H3 from mitotic chromatin in mutant lines is an indicator of H3S10p’s role in stabilising the nucleosomes and related loss of transcription in response to decreased accessibility of the promoters (Fig. 5A-B). Additionally, mitotic chromsomes retain their structure even after complete loss of condensins (Gibcus *et al*., 2018), hinting at alternate chromosome compaction mechanisms. Our data also builds on other suggestions that H3S10p allows the first degree of mitotic chromatin compaction that is then read by condensins for extreme packing of chromosomes (Giet and Glover, 2001; Hirano, 2005, 2015; Bauer, Hartl and Bosco, 2012).

### Spreading of H3S10p in mitosis around promoters

Mitotic H3S10p may have two functions; a) Marking of active domains: It spreads in mitosis and marks the active domains and promoters in proportion to their interphase transcriptional activity. The levels of H3S10p are then decoded to mimic and re-establish the marked region’s transcriptional potential from the mother to daughter cells. b) Optimal compaction and repression of active chromatin: Deficiency in mitotic H3S10p deposition at active chromatin regions results in their reduced mitotic compaction, making the region more susceptible to an enhanced transcriptional reactivation in the next interphase. Hence, marking and compaction of active domains by mitotic H3S10p seems to be crucial for the faithful propagation of a cell’s 3D-chromatin structure as well as transcription through mitosis. We speculate that spreading of H3S10p should be a pre-requisite for both of these aspects. We propose that active promoters recruit AURKB. This is followed by H3S10 catalysis and spreading of H3S10p into surrounding chromatin regions. TADs remain organized by CTCF/Cohesin complexes in the S-phase (Faseela, Notani and Sabarinathan, 2023); followed by a gradual loss of 3D chromatin architecture in G2-M transition. Aurora B Kinase activity peaks during early mitosis and probably the targeted spreading of H3S10p in the active domains/TADs helps retain their identity during mitosis. Limited kinase availability, as opposed to an enormous supply of histone H3 when coupled with either a “read-write” mechanism post loading, or “nucleosome breathing and diffusion” into proximal chromatin may bring about the spreading of mitotic H3S10p into active interphase domains, beginning at promoters. This could be one of the mechanisms in maintaining active 3D epigenetic memory through mitosis as previously described for repressive domain expansion and retention (De and Kassis, 2017; Owen, Osmanović and Mirny, 2023).

### Association of H3S10p marked islands and TAD reformation post-mitosis

Interphase 3D architecture is lost in mitosis and we have limited insight into its recovery in the next interphase. We show that highly active domains and TADs are marked by H3S10p islands in mitosis in an acetylation dependent manner and recover early post-mitosis (Fig. S3G-H). What remains unexplored is how the gradation in H3S10p levels is then decoded to restore domains/TADs upon mitotic exit. The marked domains show hyper-compaction in proportion to the H3S10p levels in mitosis (Fig. 4H-I). We hypothesize that the close proximity of regulatory sequences in this compact environment allows faster re-establishment of chromatin contacts to ensure early re-emergence of the 3D-interactions within these domains/TADs. The proximity and hence the time taken for reformation of each domain would in turn depend upon the amount of compaction and hence the levels of H3S10p in mitosis. Later into the interphase, nucleosome breathing and exchange of H3 with H3.3 might cause the dilution/removal of H3S10p from these domains, leaving behind basal levels.

## Limitations of the study

We show that mitotic compaction of active chromatin by H3S10p in mitosis is important for exact recapitulation of 3D structure and transcription in the next interphase. Our observations indicate that H3S10p spreads and forms islands in mitosis to mark the active interphase domains. However, it remains unclear how H3S10p is restricted to active domains and does not spread further into the adjacent chromatin regions. This becomes exciting to address, given the domain boundaries become weak in mitosis and the 3D architecture is lost (Naumova *et al*., 2013).

Next in our study we used heterotypic expression of histones to explore the functional relevance of H3S10p mediated mitotic compaction in gene expression in the next interphase. This is a drawback of the mammalian system we use, where mutating endogenous histones is a challenge. Complete absence of H3S10p in mitosis causes a genome-wide loss of mitotic chromatin compaction and genomic instability (Wei *et al*., 1999; Ramos-Alonso *et al*., 2023) resulting in poor survival of cells for any functional study. On the other hand, with small and transient loss of H3S10p in our study that too only in one round of mitosis, we record specific effects at active promoters in the next interphase. Positively, these observations indicate that H3S10p spreading begins at active promoters as they get affected first upon slight reduction in mitotic H3S10p.

## Materials and Methods

### Cell culture

HeLa and MCF7 cell lines obtained from the American Type Culture Collection (ATCC) were cultured in high glucose DMEM (Invitrogen) with 10% FBS (Invitrogen) and 1% Pen-Strep (Invitrogen) at 37℃ and 5% CO2 in a humidified incubator. The cells were passaged every 3-4 days.

### Cell synchronization

Mitotic vs Asynchronous cell synchronization (for ChIP seq): Cells were treated with 100 ng per ml Nocodazole (Sigma M1404) for 24 hours with DMSO (Sigma) as the solvent control (Naumova *et al*., 2013; Ma and Poon, 2017; Surani *et al*., 2021).

Mitotic cell synchronization (for stable lines experiments): Cells were first arrested in early S with a single Thymidine (2mM; Sigma T1895) block for 24h followed by a release into fresh media for 8h with or without induction. Subsequently, Nocodazole was added for 12h, and cells were harvested by shake-off/trypsinization. Doxycycline (1mg/ml; Sigma D9891) was used for induction.

Next interphase (for stable lines experiments): Mitotic cells obtained from the above method were washed with 1X-DPBS (Invitrogen) and twice with DMEM, following mitotic shake-off. Cells were then plated in fresh media and allowed to grow for 4-13h in the next interphase before harvesting.

### Cloning of H3S10 mutant plasmids

Human histone H3 gene with an N terminus HA tag (HA: H3.1 429bp) was cloned in pCW- Cas9, which was a kind gift from Eric Lander & David Sabatini (Addgene plasmid #50661; http://n2t.net/addgene:50661; RRID:Addgene_50661) (Wang *et al*., 2014), after removing Cas9 with Restriction Endonucleases: NheI (NEB R0131L) and BamHI-HF (NEB R3136L). Three variants of the H3 gene with HA tag; namely, HA-H3S10 (WT), HA-H3S10A (phospho-dead mutant), and HA-H3S10D (phospho-mimetic mutant) were PCR amplified in two rounds: a) from HeLa genomic-DNA with common HA-H3 Forward and H3 Reverse oligos; b) the obtained PCR product was further amplified with respective variant HA-H3 Forward and common H3 reverse oligos (see Supplementary Table 1; Oligos). The inserts were digested and ligated into the digested vector backbone. The plasmids were verified by Sanger sequencing. HA-H3 expression was under the control of doxycycline.

### Lentiviral transduction

HEK293FT cells were seeded in poly-D-Lysine coated dishes. Cells were transfected with the plasmid of interest (mentioned in ‘Cloning of H3S10 variant plasmids’) and the lentiviral packaging plasmid pCMV-VSV-G a gift from Bob Weinberg (Addgene plasmid #8454; http://n2t.net/addgene:8454; RRID:Addgene_8454) (Stewart *et al*., 2003) and PAX2 using Lipofectamine 2000 and media was changed after 6 hours. Viral soup collected from 48 and 72 hours was pooled together, filtered, and added to HeLa cells along with 8ug/ml of polybrene. Infection was terminated after 16 hours of transduction.

### Generation of HeLa HA-tagged H3 stable lines

HeLa cells transduced with either of the three H3 histone variant plasmids (H3S10, H3S10A, and H3S10D) were put under 5ug/ml Puromycin (Gibco, A11138-03) selection. Media was changed every third day with antibiotics replenished. The lines were tested for induction of the HA-tagged H3 histones with Doxycycline treatment for 24 hours (Fig. 4A). Mutations in the stable line were confirmed by sanger sequencing of the genomic DNA and HA- Immunoprecipitaion assay (Fig. S4C-D).

### Antibodies

H3S10p (ab5176, ab14955), HA (ab9110, CST 3724), H3 (ab1791), GAPDH (sc32233).

### Immunostaining

Cells were grown on coverslips or plated on coverslips (for the next interphase experiments after mitotic shake-off) and washed with 1X-PBS (X2) following harvest. Fixation was done using 4% PFA (Sigma) for 12’ at RT, followed by three 1X-PBS washes and permeabilization with 0.5% Triton-X100 in 1% BSA for 15’ at RT. Cells were then incubated with primary antibody overnight at 4°C and were given three 1XPBST washes. Fluorophore secondary antibody incubation was done for 1h at RT followed by 1X-PBST (X3) washes. Coverslips were then mounted using 90% glycerol after 15’ incubation (RT) with Hoechst 33342. For comparing bound/soluble fractions; 0.5ml CSK buffer (10mM PIPES/KOH pH 6.8, 100mM NaCl, 300mM sucrose, 1mM EGTA, 1mM MgCl2; with fresh addition of 1mM DTT, 1X PIC, and 0.5% Triton-X) per well was used before fixation.

### Metaphase spread

Metaphase spreads were prepared as described in (Naumova *et al*., 2013). Briefly, mitotic cells were harvested at 1600g at RT for 5’. The cell pellet was then resuspended in 0.75M KCl buffer and kept at 37°C for 20’. After a brief spin at 1600g, RT; the swollen pellet was carefully resuspended in 3:1 Methanol: Glacial acetic acid (fixative solution) with drop-by-drop addition. The sample was then stored at 4°C. For staining, the cells were pelleted and resuspended in fresh fixative every time and dropped on a coverslip from a 30cm height for the mitotic nuclei to burst. The coverslips were then dried and proceeded for immunostaining as mentioned above.

### Microscope imaging and analysis

Slides were imaged at Olympus FV3000 (PLAPON) 60x/1.42 oil objective. The median stack was used for making line ROIs and PlotProfile tool was used to plot the ROIs for different stages. A custom-generated MATLAB script was used for intensity calculation. Hoechst 33342 channel was used to generate a 3D nuclear mask. Signal intensity within the 3D nuclear mask for different channels was calculated. For metaphase spread area calculation, binary mask was created after thresholding for 30 metaphase spreads in each condition and area covered by chromosomes in each nuclear ROI was measured in Fiji (Schindelin *et al*., 2012).

### Immunoprecipitation

For IP, freshly procured cell pellet from ∼3mn cells was resuspended in 200ul of HLB (hypotonic lysis buffer; 10mM Tris pH7.5, 10mM NaCl, 3mM MgCl2, 0.3%NP40, 10% Glycerol, 1X PIC, 1mM NaF) and kept on ice for 10’. The cells were then spun at 4000 rpm, 4℃ for 5’ and the cytosolic fraction was discarded as the supernatant. The nuclear pellet was resuspended in 250ul NP40 Buffer (1% NP40, 150mM NaCl, 50mM Tris pH8, 1mM EDTA, 0.1% SDS, 1mM DTT, 1X PIC, 1mM NaF) and kept on ice for 40’. The nuclear fraction was then sonicated for 10 cycles (30” ON/30” OFF) and spun at 12000 rpm for 12’ at 4℃ to remove debris. The soup was made up to 2.1X parts (total 1100ul) such that 2 equal parts, 500ul each were used for IP (with HA antibody) and bead while 0.1 fraction was kept as input at 4℃ for 4h on the rotor. Pre-blocked Protein G Dynabeads were added to the lysates for 1h at RT. The beads were washed thrice with NP40 Buffer (300mM NaCl total) and 2X laemmli dye was added to the beads while 6X laemmli dye was added to the input and incubated at 95℃ for 10’.

### Fractionation

Cells were harvested by mitotic shake-off, washed with cold 1X-PBS, and pelleted at 200g, 4℃. 0.2ml of cold CSK buffer (recipe same as above in immunostaining) was then added per million cells pellet and pipetted. The tubes were kept on ice for 20’ with gentle agitation by tapping in between. This was followed by a spin at 800g, 4℃. The supernatant obtained was kept as the soluble fraction while the pellet was the bound fraction, with 6X and 2X laemmeli dye addition respectively.

### Immunoblotting

Total lysates were obtained by adding 2X laemmeli dye directly to the cell pellets. The protein lysate samples were run in a 15% SDS-PAGE gel (Biorad), followed by protein transfer (Invitrogen) in Tris-glycine buffer at 25V (90’) on ice using PVDF membrane (Millipore). The membrane was blocked in 5% BSA or skimmed milk for 1h at RT (except for H3, H3S10p: overnight blocking at 4℃), followed by primary antibody incubation overnight at 4℃ (except H3 and H3S10p which were kept for 4h). Blots were then washed before and after HRP secondary antibody incubation (1h RT) with 1X-TBST. The signal was amplified with ECL substrate (GE Healthcare, RPN2106) and detected in Image Quant LAS4000 using a CCD camera.

### FACS

Cells were harvested and fixed with 80% cold ethanol drop by drop on ice (20’). The cells were then pelleted and stored at -20℃. Cells were stained with cold PI buffer (0.1% Sodium citrate, 25ug/ml Propidium Iodide, 0.03% NP40, 40ug/ml RNaseA) before subjecting them to BD FACS Fortessa for cell cycle profiling. The DNA content assessment was done using 488 laser line. The cell cycle profile images were obtained from the system software.

### Chromatin Immunoprecipitation (ChIP) and ChIP-seq

For ChIP, 10 million cells were cross-linked using 1% formaldehyde (Sigma) for 10’ at RT, and the reaction was quenched with 125 mM glycine for 5’ at RT. The cells were washed and scraped in cold PBS on ice. After spinning the samples at 2000 rpm, 5’ and 4°C; the cell pellets were then stored at -80°C until further use. The pellets were thawed on ice and lysis was done with ∼0.8ml NLB (Nuclear lysis Buffer; 50mM Tris-HCl pH 7.4, 1% SDS, 10mM EDTA pH 8.0, and 1X PIC with 10mM NaF) on ice for 10’. The lysed fraction was sheared in a water bath sonicator (Diagenode Bioruptor- 28 cycles: 30” ON/30” OFF and Covaris- 20’ at 175 Peak power, 10 Duty factor and 200 cycles/Burst) to generate chromatin fragments (400-500 bp in length). Chromatin lysate was cleared by centrifugation at 12000 rpm, 4°C for 12 min. A total of 100ug (65ug for H3S10p) chromatin was taken per IP and Bead sample and diluted with DB (Dilution Buffer; 20mM Tris-HCl pH 7.4, 100mMNaCl, 2mM EDTA pH 8.0, 6510.5% Triton X-10, and 1X PIC with 10mM NaF) 2.5 times. 10% of this was taken as input DNA. The IP samples with antibody (1ug) and input, beads without antibody, were rotated at 4°C overnight. 15ul pre-blocked (in 1% BSA, 1h, 4°C) Protein G Dynabeads were incubated with the antibody-bound and beads lysate for 4h on rotor at 4°C. The bead samples were then washed sequentially with Wash Buffer I (20 mM Tris-HCl pH 7.4, 150 mM NaCl, 0.1% SDS, 2 mM EDTA pH 8.0, 1% Triton X-100), Wash Buffer II (20 mM Tris-HCl pH 7.4, 500 mM NaCl, 2 mM EDTA pH 8.0, 1% Triton X-100), and Wash Buffer III (10 mM Tris-HCl pH 7.4, 250 mM LiCl, 1% NP-40, 1% Sodium Deoxycholate, 1 mM EDTA pH 8.0; followed by a 1X TE (10 mM Tris pH 8.0, 1mM EDTA pH 8.0) wash. The chromatin complexes were then eluted using 200ul EB (Elution Buffer; 100mM NaHCO3, 1% SDS) at 37°C, 30’-90’ in a thermomixer at 1400rpm. All samples were reverse cross-linked at 65°C for 8–16 h with 14ul of 5M NaCl. DNA was purified by phenol-chloroform extraction followed by ethanol precipitation and the DNA pellet obtained was dissolved in 100ul 1X TE (for hand qPCRs) or 10ul NFW (Nuclease Free Water; for sequencing). Library was prepared using the NEB Next ChIP-seq library preparation kit (E7103L) as per manufacturer instructions. The samples were sequenced on HiSeq 2500 (Illumina) and NovaSeq 6000 (Illumina) platforms using a 1x50bp and 2X50bp sequencing format respectively at the NCBS NGS Facility.

### ChIP-seq analyses

The sequenced reads were checked for QC using FASTQC and aligned to hg19 using default Bowtie2 (Langmead and Salzberg, 2012) options. The BAM files were read depth normalized using the SAM tool and were subjected to MACS2 (with default options including broad peak calling, with input as control) to get bedGraph files, which were then converted to bigwigs using wig-to-bedGraph for visualization. These tracks were uploaded to the UCSC genome browser and shared as processed files. The BAM/Bigwig files were used for multibam/multibigwig summary tools to get normalized coverage values. Computematrix and plotHeatmap were used for calculating and plotting the signal intensity at regions of interest. The analysis was performed on the Galaxy platform (Afgan *et al*., 2018; The Galaxy Community, 2022). Publicly available bigwig files for different histone modifications and PolII were downloaded from ENCODE or NCBI (see Supplementary Table 2; NGS datasets) (‘An Integrated Encyclopedia of DNA Elements in the Human Genome’, 2012; Luo *et al*., 2020) and used directly for plotting heatmaps and calculating enrichment. ChIP seeker was used for genome annotation. IGV Desktop application was used to visualize data (Robinson *et al*., 2011).

### RNA isolation and RNA-seq

RNA was isolated using Trizol (Invitrogen) as per the supplier’s manual. Library was prepared using NEBNext RNA depletion and NEBNext Ultra II Directional RNA Library Prep Kits as per the instructions. The samples were sequenced on HiSeq 2500 (Illumina) and NovaSeq 6000 (Illumina) platforms using a 1x50bp and 2x50bp sequencing format respectively in the NCBS NGS Facility.

### RNA-seq analysis

The raw fastq files were subjected to Trimmomatic and FASTQC was done to check read quality. The reads were then aligned to hg19 using HISAT2. BAM files obtained were read depth normalised and FeatureCounts was used with default settings to get coverage at all GeneIDs. The fold changes between different lines were calculated using DeSeq2. The analysis was performed on the Galaxy platform (Afgan *et al*., 2018; The Galaxy Community, 2022). For making gene categories based on their expression HeLa asynchronous RNA seq data (ENCFF622JEX) was downloaded from ENCODE. IGV Desktop application was used to visualize data (Robinson *et al*., 2011).

### HiC analysis

TADs called from asynchronous HeLa population using HOMER were used (Islam *et al*., 2023). Processed HiC files (.hic) for mitotic release (Abramo *et al*., 2019); were downloaded from the 4DN portal (Dekker *et al*., 2017; Reiff *et al*., 2022) (see Supplementary Table 2; NGS datasets). The .hic files were converted to (.cool) format using hic2cool tool, for making pile-up plots using coolpup.py and plotpup.py. FastQ files for mitotic release (Abramo *et al*., 2019) were downloaded from EBI SRA and HiC analyses were performed using HOMER (Heinz *et al*., 2010) with default settings to obtain A/B compartments and PC1 file. The PC1 score at 25 kb resolution, from different time points after mitosis was used to measure the strength of A/B compartments from the asynchronous stage. TAD boundary strength in the asynchronous population was calculated using insulation score file.

### GRO-seq analysis

Publicly available files were downloaded (see Supplementary Table 2; NGS datasets), and used to plot signal and calculate enrichment at regions of interest using deep tools.

### ATAC-seq

ATAC samples were prepared following the protocol described in (Dhall *et al*., 2023). Briefly, cells were lysed using lysis buffer, and single nuclei were isolated. The nuclei were tested for viability using DAPI and 50,000 nuclei were taken for tagmentation reaction as per the manual (Illumina tagmentation kit). The processed samples were subjected to library preparation similar to ChIP-seq methods.

### ATAC-seq analyses

The data was analyzed using default parameters as per the Galaxy ATAC-seq analysis guidelines (https://training.galaxyproject.org/training-material/topics/epigenetics/tutorials/atac-seq/tutorial.html, Hiltemann *et al.,* 2023). The processed bigwig files were then used to plot the signal at published ATAC peaks from interphase and mitosis (Shen *et al.,* 2022) in HeLa. IGV Desktop application was used to visualize data (Robinson *et al*., 2011).

### Allele specific analysis

GATK ASEReadCounter (Auwera and O’Connor, 2020) (v4.1.9.0) was used for allele specific analysis. Reads with minimum base quality of 10 and minimum read mapping quality of 20 were considered for the analysis. Only those heterozygous SNPs with minimum read depth of 8 (for gene expression) and 5 (for ChIP-Seq) were considered. At feature level, cutoff of 15 reads at heterozygous SNP positions in total was used. WGS read depth at heterozygous SNPs was used for copy number correction. To determine the significance of allele specific activity with respect to copy number status at each feature level from haplotype A, binomial test followed by bonferroni correction was done.

RNA-seq data and phased heterozygous SNP positions from HeLa genome were obtained from the controlled access dbGaP repository phs000640.v1.p1.

### Gene expression analysis in cancers

TCGA-CESC (Cervical squamous cell carcinoma and endocervical adenocarcinoma, n=301)(Burk *et al*., 2017) gene expression data was obtained from GDC portal (https://portal.gdc.cancer.gov/).

## Contributions

SM and DN conceived the project. SM performed most experiments with help from AN, AKS, and RM. SM and SM (Mishra) performed informatics analysis. SM and DN wrote the manuscript with input from all the authors. All authors read the manuscript.

## Acknowledgments

Authors acknowledge the critical inputs from Drs. Sachin Kotak, Ravi Muddashetty, and Rajesh K. Ladher as Thesis committee members to SM. We thank Dr. Srimonta Gayen and Avinchal Manhas for providing mESCs and helping with their culture. We are grateful to NCBS-CIFF (Central Imaging and Flow cytometry Facility) and NGS (Next Generation Sequencing) Facility for their services. We acknowledge the intra-mural funding from TIFR-NCBS (12-R&D-TFR-5.04-0800). The work was funded by fellowship from DBT-Wellcome Trust IA (IA/1/14/2/501539) and EMBO Global Investigator award to DN. SM, AKS and RM are supported by the PhD program of TIFR-NCBS. AN acknowledges fellowship from DBT/CSIR. SM (Mishra) is funded by the UGC JRF. We thank Umer Farooq for help with stable lines, and Bharath Saravanan and Deepanshu Soota for their critical inputs. We thank all DN lab members for constructive discussions.

## Data Availability

The data can be accessed on accession code GSE252349.

---

**Figure S1.**
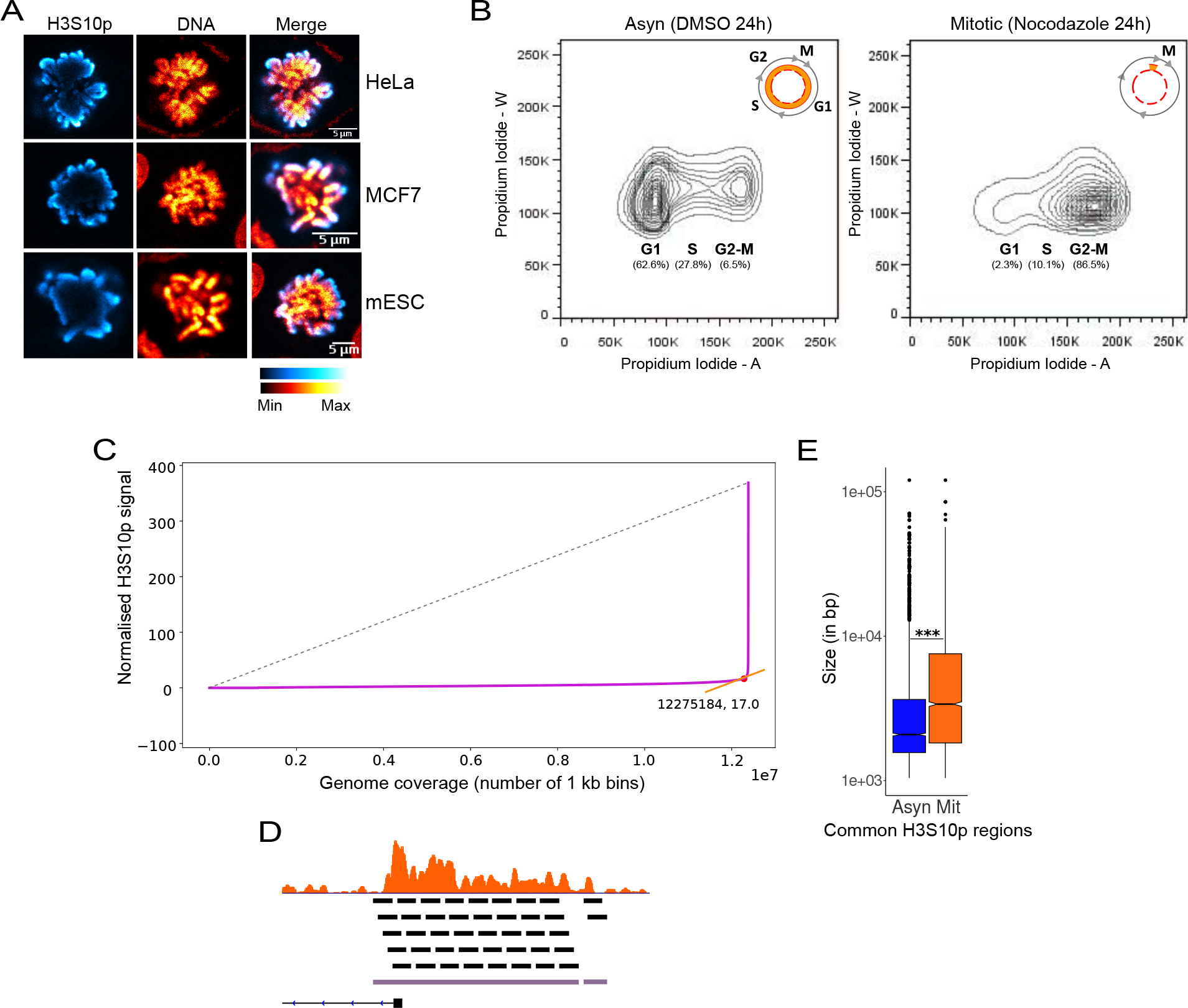
Related to Figure 1 A. Immunostained prometaphase cells from three cell lines showing H3S10p (pseudocoloured, Cyan Hot) and DNA stained with Hoechst (pseudocoloured, Red hot). B. FACS profile of the Asyn and Mit samples showing the shift in DNA (Stained with Propidium Iodide) from G1-S-G2M to G2-M. C. Graph showing the threshold used for selecting 1kb windows of H3S10p enrichment for downstream processing (See methods), indicated by the interception of the curve with H3S10p signal on the y-axis and genomic coverage on the x-axis. D. Schematic for the stitching of sliding 1kb windows (in black) to obtain a region (in violet) of H3S10p enrichment (orange). E. Boxplot showing the size distribution of common H3S10p regions in asynchronous and mitotic populations. Statistics details: p-value >0.05 (ns), <=0.05 (*), <=0.01 (**), and <0.001 (***). Statistical significance was determined by the Wilcoxon signed-rank test (paired).

**Figure S2.**
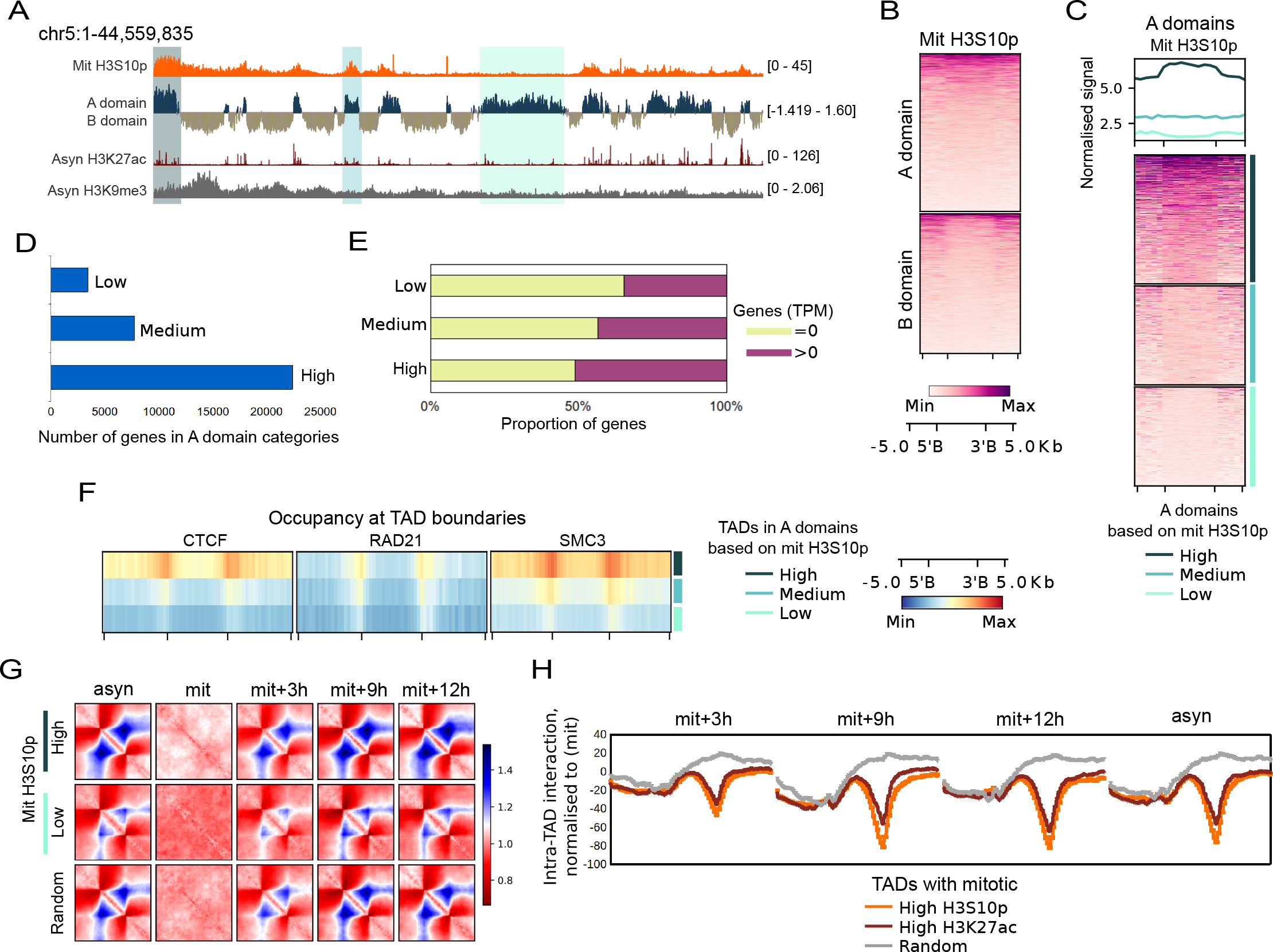
Related to Figure 2. A. Asynchronous compartments (super domains- A and B) showing H3S10p enrichment restricted within A domain boundaries, varying with asyn H3K27ac levels and antagonising with asyn H3K9me3 levels. B. Heatmap showing Mit H3S10p signal in A and B superdomains. C. Heatmap showing Mit H3S10p signal in the three A domain categories divided based on mitotic H3S10p signal in them. D. Barplot showing the number of genes in the three A domain categories. E. Barplot showing the proportion of genes in Fig. S5C, with expression in interphase. F. Profile plot showing CTCF, RAD21, and SMC3 occupancy at TAD boundaries of the three groups. G. APA plot for local rescaled pileups of TADs with mitotic H3S10p (high and low), and random TADs, plotted using insulation score from HiC data performed at various times post mitotic release. H. Line plot for quantification of TAD insulation in high mitotic H3K27ac, H3S10p, and random TADs from the APA plot in Fig. 2H normalized to mitotic values for high H3K27ac and H3S10p TADs.

**Figure S3.**
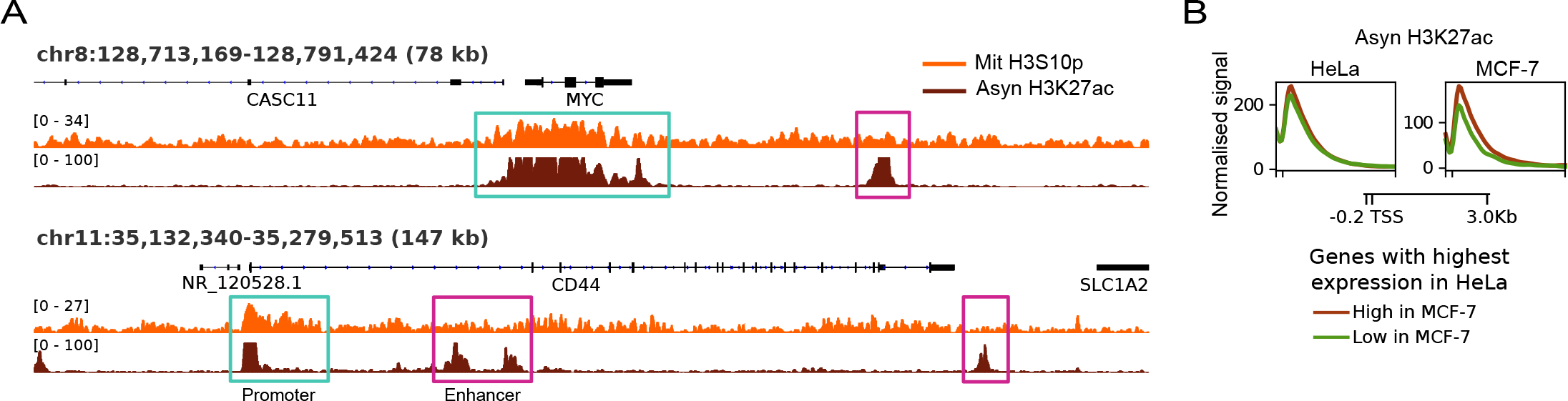
Related to Figure 3 A. IGV snapshot around MYC and CD44 gene showing high H3S10p enrichment at the gene promoters but not enhancers. B. Asyn H3K27ac levels from HeLa and MCF-7 at the two subgroups from Fig. 3H. Statistics details: p-value >0.05 (ns), <=0.05 (*), <=0.01 (**), and <0.001 (***). The Mann-Whitney U-test determined statistical significance.

**Figure S4.**
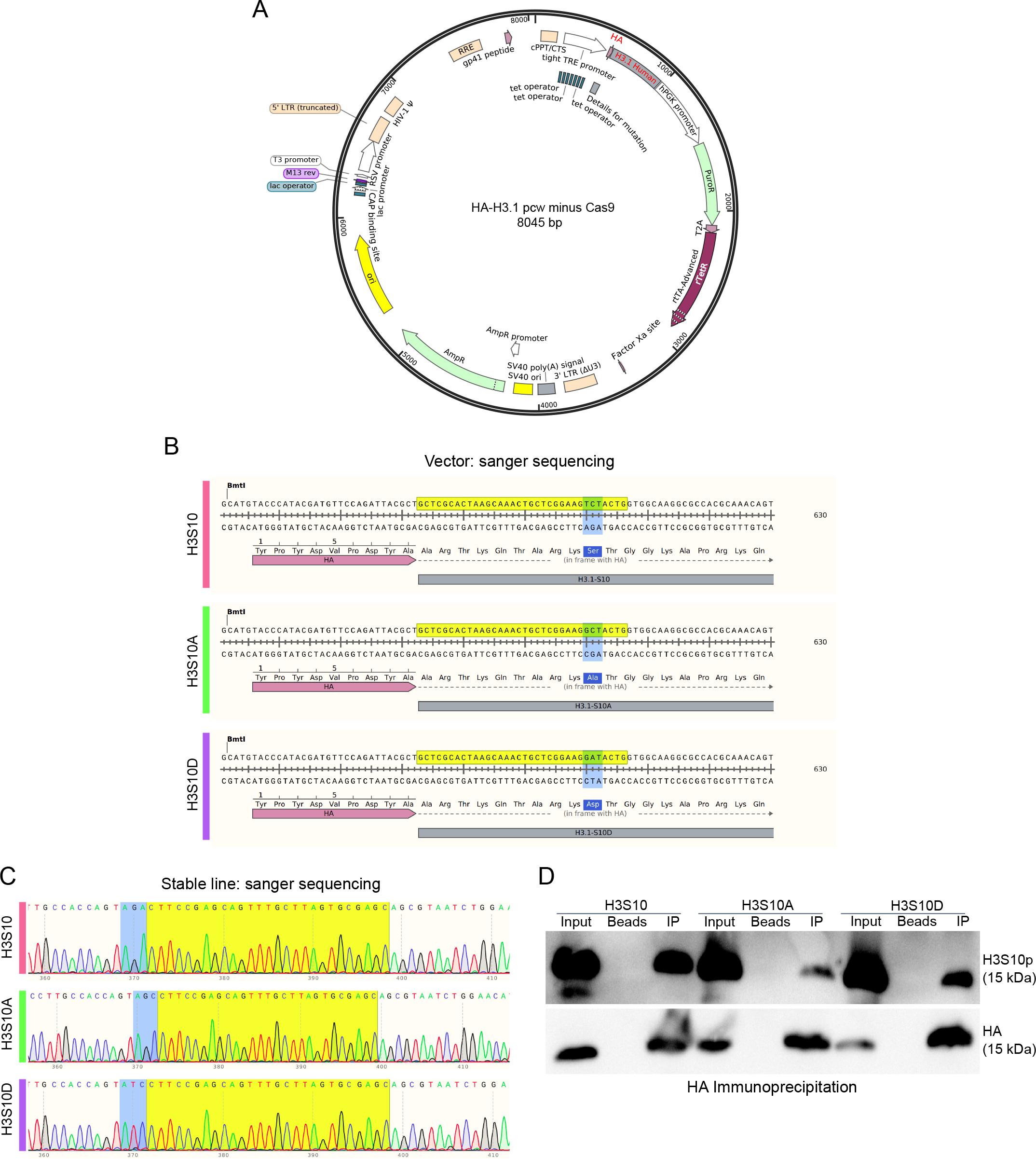
Making of stable HA-tagged histone H3 variant cell lines A. Vector map of the plasmid used for cloning HA-tagged histone H3S10 and its mutants. B. Sanger sequencing of the histone fragments from respective vectors. C. Sanger sequencing of the stable lines with HA forward and Histone H3 reverse oligos. ID Western blot for H3S10p from mutant lines, pulled down using anti-HA antibody.

**Figure S5.**
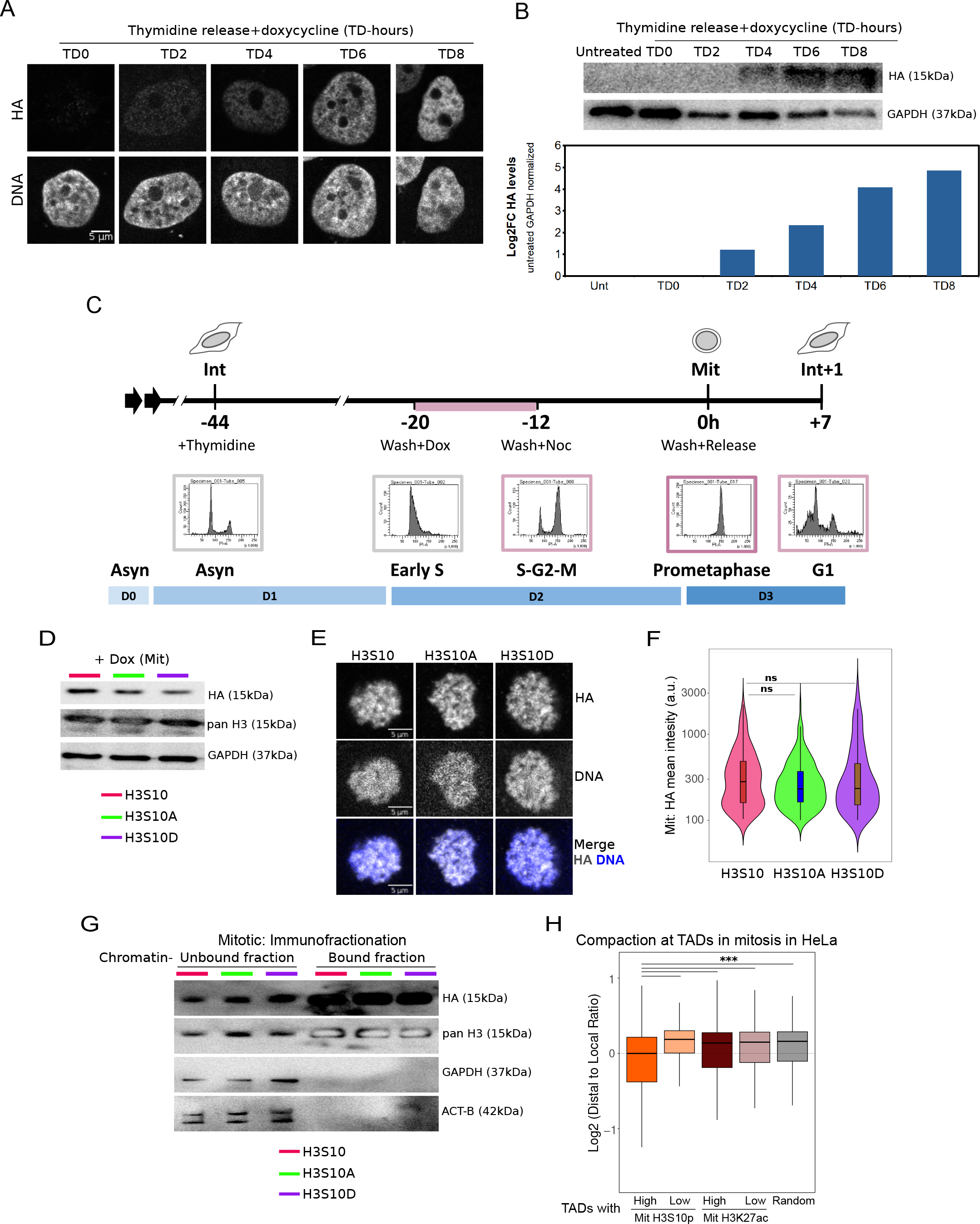
Related to Figure 4 A. Immunostainings showing HA-H3 incorporation at different time points upon doxycycline treatment, following release from early S block with thymidine. B. Immunoblots showing elevation in levels of HA-H3 with increasing doxycycline induction time after thymidine release; confirming S5A at global scale. C. Cell cycle profile from the treatment timeline from Fig. 4B. D. Western blot showing low levels of HA-tagged histone H3 variants as opposed to the total H3 levels. E. HA and H3S10p immunostaining of mitotic cells from the stable lines showing incorporation of HA-tagged histone H3 variants and characteristic mitotic H3S10p pattern. F. Quantification of HA from Fig. S5B. G. Western blot with beta-actin, GAPDH, pan H3 and HA antibodies in chromatin-bound and unbound fraction. H. Boxplots showing actual mitotic chromatin compaction measured as Log2DLR from mitotic HiC, at various TAD categories from Fig. 4I. Statistics details: p-value >0.05 (ns), <=0.05 (*), <=0.01 (**), and <0.001 (***). The Mann-Whitney U-test determined statistical significance.

**Figure S6.**
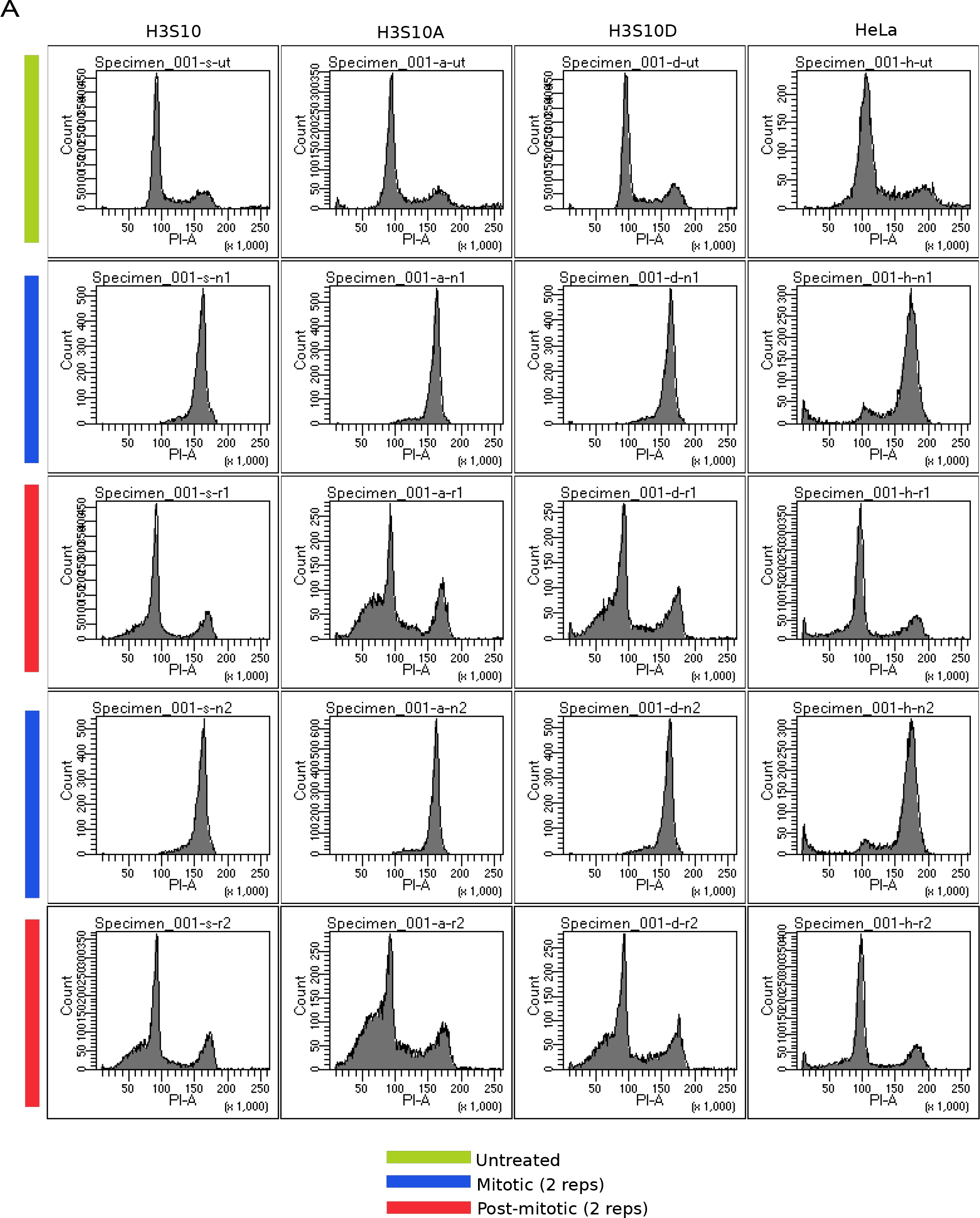
Mitotic and post-mitotic cell cycle profile of the stable lines A. Cell cycle FACS profile of the three stable lines: H3S10, H3S10A, and H3S10D harvested at different time points of the treatment regime mentioned in Fig. 4B.

**Figure S7.**
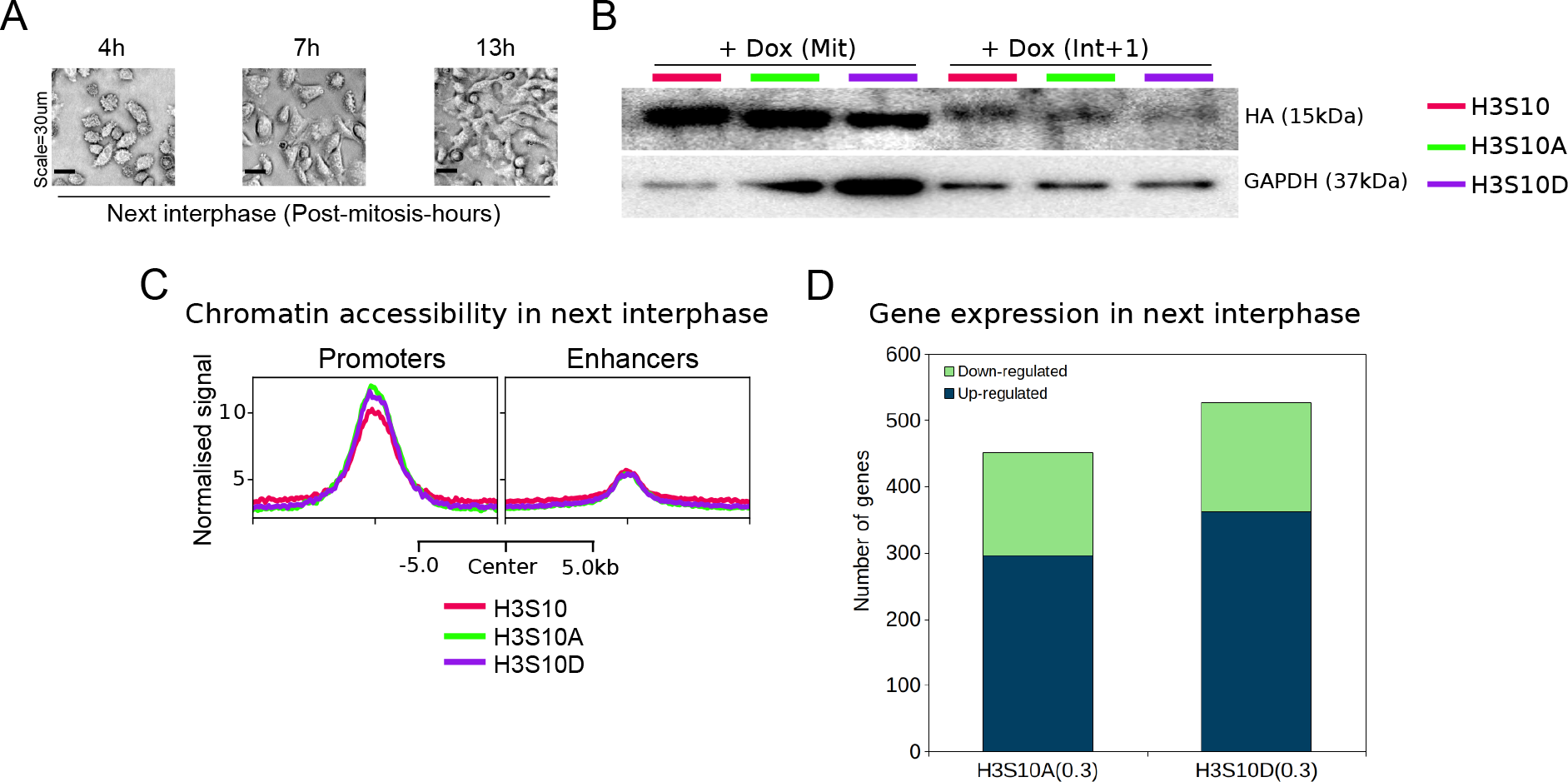
Related to Figure 5 A. DIC images of HeLa cells at different time points post mitotic release. B. Western blot showing non-detectable levels of HA-tagged H3 histones in the next interphase compared to the mitotic levels as a result of mitosis-specific induction. C. Profile line plot showing post-mitotic chromatin opening at promoters and enhancers in mutant lines. D. Number of dysregulated genes in mutant lines showing more upregulation than downregulation. Statistics details: p-value >0.05 (ns), <=0.05 (*), <=0.01 (**), and <0.001 (***). The Mann-Whitney U-test determined statistical significance.

**Supplementary Table 1:**
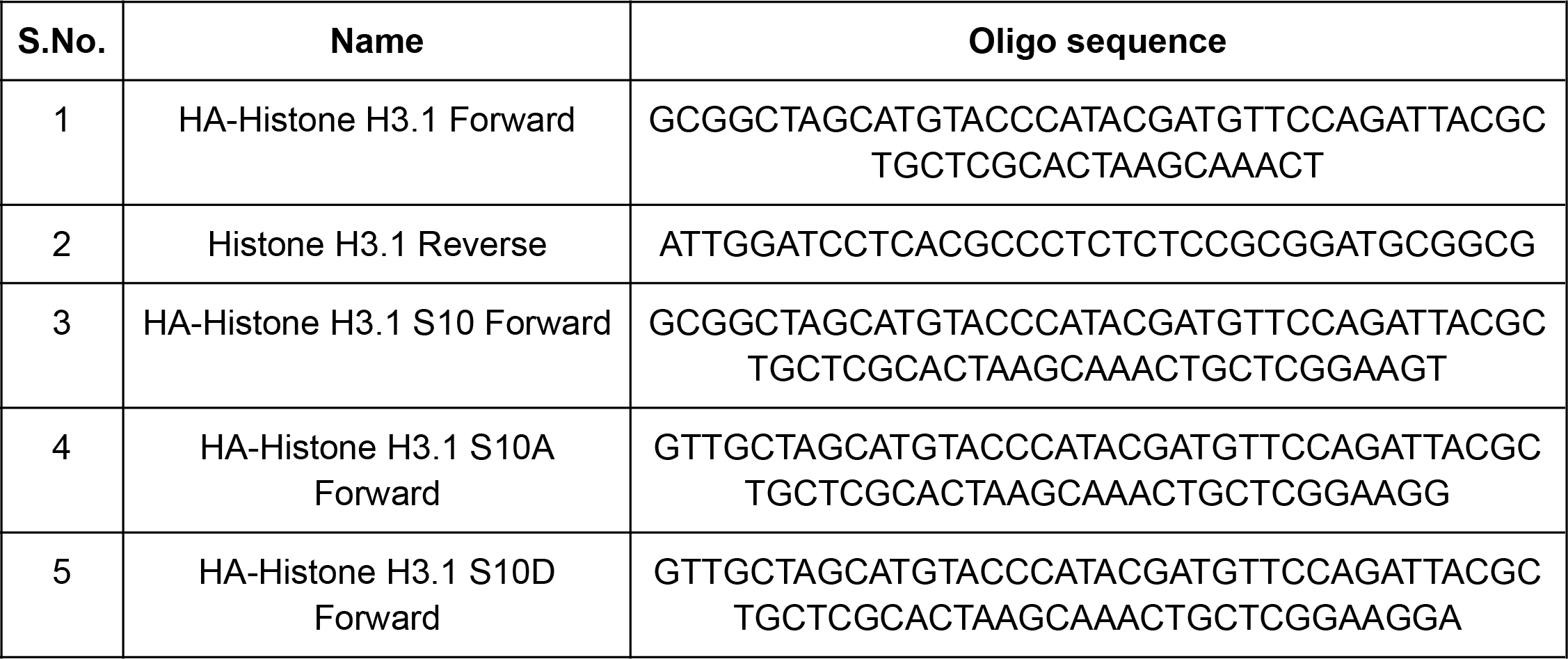
Oligos.

**Supplementary Table 2:** NGS datasets.

| GEO |  |  |  |  |
| --- | --- | --- | --- | --- |
| S.no. | Accession # | Experiment type, cell type | Experiment detail | Reference |
| 1 | GSM2891724 | H3K27ac ChIP-seq, HeLa-S3 | Untreated (asynchronous) | (Javasky *et al.*, 2018) |
| 2 | GSM2891722 | H3K27ac ChIP-seq, HeLa-S3 | Synchronized (mitotic) |  |
| 3 | GSM5133464 | H3K9ac ChIP-seq, HeLa-S3 | Untreated (asynchronous) | (Gandhi *et al.*, 2022) |
| 4 | GSM5133466 | H3K9ac ChIP-seq, HeLa-S3 | Synchronized (mitotic) |  |
| 5 | GSM5848498 | ATAC-seq, HeLa-S3 | Interphase | (Shen *et al.*, 2022) |
| 6 | GSM5848500 | ATAC-seq, HeLa-S3 | Mitosis |  |
| 7 | GSM1846808 | Nascent RNA-seq, HeLa | Asynchronous | (Liang *et al.*, 2015) |
| 8 | GSM1846809 | Nascent RNA-seq, HeLa | Mitotic |  |
| 9 | GSM6123347 | RNA-seq, MCF-7 | Untreated (asynchronous) | (Xu *et al.*, 2023) |
| 10 | GSM3909704 | in situ Hi-C, HeLa-S3 | Prometaphase arrest (mitotic) | (Abramo *et al.*, 2019) |
| 11 | GSM3909694 | in situ Hi-C, HeLa-S3 | 3 hours release after prometaphase arrest |  |
| 12 | GSM3909688 | in situ Hi-C, HeLa-S3 | 6 hours release after prometaphase arrest |  |
| 13 | GSM3909685 | in situ Hi-C, HeLa-S3 | 9 hours release after prometaphase arrest |  |
| 14 | GSM3909722 | in situ Hi-C, HeLa-S3 | Before prometaphase arrest (asynchronous) |  |
| 15 | GSE252349 | ChIP-seq_H3S10p_Asyn_Repeat_1 | Asynchronous | This Study |
| 16 |  | ChIP-seq_H3S10p_Asyn_Repeat_2 | Asynchronous |  |
| 17 |  | ChIP-seq_H3S10p_Asyn_Repeat_3 | Asynchronous |  |
| 18 |  | ChIP-seq_H3S10p_Mit_Repeat_1 | Mitotic |  |
| 19 |  | ChIP-seq_H3S10p_Mit_Repeat_2 | Mitotic |  |
| 20 |  | ChIP-seq_H3S10p_Mit_Repeat_3 | Mitotic |  |
| 21 |  | ChIP-seq_input_Asyn | Asynchronous |  |
| 22 |  | ChIP-seq_input_Mit | Mitotic |  |
| 23 |  | ChIP-seq_MCF-7_H3S10p_Asyn | Asynchronous |  |
| 24 |  | ChIP-seq_MCF-7_H3S10p_Mit | Mitotic |  |
| 25 |  | ChIP-seq_HA_-S10_Mit_Repeat_1 | 7 hours post-mitotic |  |
| 26 |  | ChIP-seq_HA_-S10A_Mit_Repeat_1 | 7 hours post-mitotic |  |
| 27 |  | ChIP-seq_HA_-S10D_Mit_Repeat_1 | 7 hours post-mitotic |  |
| 28 |  | ChIP-seq_H3S10p_-S10_Mit_Repeat_1 | 7 hours post-mitotic |  |
| 29 |  | ChIP-seq_H3S10p_-S10A_Mit_Repeat_1 | 7 hours post-mitotic |  |
| 30 |  | ChIP-seq_H3S10p_-S10D_Mit_Repeat_1 | 7 hours post-mitotic |  |
| 31 |  | ChIP-seq_H3_-S10_Mit_Repeat_1 | 7 hours post-mitotic |  |
| 32 |  | ChIP-seq_H3_-S10A_Mit_Repeat_1 | 7 hours post-mitotic |  |
| 33 |  | ChIP-seq_H3_-S10D_Mit_Repeat_1 | 7 hours post-mitotic |  |
| 34 |  | ATAC-seq_S10_Mit_7h_Repeat_1 | 7 hours post-mitotic |  |
| 35 |  | ATAC-seq_S10A_Mit_7h_Repeat_1 | 7 hours post-mitotic |  |
| 36 |  | ATAC-seq_S10D_Mit_7h_Repeat_1 | 7 hours post-mitotic |  |
| 37 |  | RNA-seq_S10_Mit_7h_Repeat_1 | 7 hours post-mitotic |  |
| 38 |  | RNA-seq_S10_Mit_7h_Repeat_2 | 7 hours post-mitotic |  |
| 39 |  | RNA-seq_S10A_Mit_7h_Repeat_1 | 7 hours post-mitotic |  |
| 40 |  | RNA-seq_S10A_Mit_7h_Repeat_2 | 7 hours post-mitotic |  |
| 41 |  | RNA-seq_S10D_Mit_7h_Repeat_1 | 7 hours post-mitotic |  |
| 42 |  | RNA-seq_S10D_Mit_7h_Repeat_2 | 7 hours post-mitotic |  |

4D Nucleome Data Portal
| S.No. | Accession # | Experiment | Experiment detail | Reference |
| --- | --- | --- | --- | --- |
| 1 | 4DNFIMF6CFOM | in situ Hi-C, HeLa-S3 | Prometaphase arrest (mitotic) | (Abramo *et al.*, 2019) |
| 2 | 4DNFIDT9EB5M | in situ Hi-C, HeLa-S3 | 3 hours release after prometaphase arrest |  |
| 3 | 4DNFIVOJGWNP | in situ Hi-C, HeLa-S3 | 6 hours release after prometaphase arrest |  |
| 4 | 4DNFI9ZBEBJH | in situ Hi-C, HeLa-S3 | 9 hours release after prometaphase arrest |  |

| 5 | 4DNFIO8HVKOL | in situ Hi-C, HeLa-S3 | Before prometaphase arrest (asynchronous) |  |
| --- | --- | --- | --- | --- |
| **ENCODE** |  |  |  |  |
| **S.No.** | **Accession #** | **Experiment** | **Experiment detail** | **Reference** |
| 1 | ENCSR000AOC | H3K27ac ChIP-seq, HeLa-S3 | Untreated (asynchronous) | ENCODE portal, https://www.encodeproject.org/ |
| 2 | ENCSR340WQU | H3K4me3 ChIP-seq, HeLa-S3 | Untreated (asynchronous) |  |
| 3 | ENCSR000ECT | Pol-II S2p ChIP-seq, HeLa-S3 | Untreated (asynchronous) |  |
| 4 | ENCSR000EDE | RAD21 ChIP-seq, HeLa-S3 | Untreated (asynchronous) |  |
| 5 | ENCSR000ECS | SMC3 ChIP-seq, HeLa-S3 | Untreated (asynchronous) |  |
| 6 | ENCSR000AOA | CTCF ChIP-seq, HeLa-S3 | Untreated (asynchronous) |  |
| 7 | ENCSR752UOD | H3K27ac ChIP-seq, MCF-7 | Untreated (asynchronous) |  |
| 8 | ENCSR000CPR | polA plus RNA-seq | Untreated (asynchronous) |  |
| 9 | ENCSR056UBA | H3K9ac ChIP-seq, MCF-7 | Untreated (asynchronous) |  |
| 10 | ENCSR000AOH | H3K9ac ChIP-seq, HeLa-S3 | Untreated (asynchronous) |  |

